# Regulation of motor neuron differentiation in the *Ciona* larva

**DOI:** 10.64898/2025.12.17.694982

**Authors:** Sydney Popsuj, Tenzin Kalsang, Christina D. Cota, Alberto Stolfi

## Abstract

Here we investigate the gene regulatory networks responsible for the differentiation of cholinergic neurons in the *Ciona* larval motor ganglion, which functions as the organism’s central pattern generator for swimming and dispersal. We demonstrate conserved roles motor neuron-enriched transcription factors Neurogenin and Onecut, with Neurogenin likely sitting atop the regulatory cascade. We also identify a key role for the transcription factor Nkx6 in specifying one motor neuron subtype, Motor Neuron 1 (MN1), and show that the secreted Wnt pathway inhibitor Dkk3 is required by MN1 for its unique neuromuscular endplates. We propose that Dkk3 interacts with LRP receptors in target muscle cells, through conserved NxI/V/F motifs shared with another key secreted neuromuscular synapse effector, Agrin. Taken together, our results provide critical insights into the development and evolution of cholinergic neurons involved in chordate locomotion.

## Introduction

Cholinergic neurons, those that use acetylcholine as their major neurotransmitter, have diverse functions in vertebrates ranging from processing sensory information to associative learning and neuromodulation (Ahmed et al., 2019; Picciotto et al., 2012). The neurons play a particularly critical role in providing the primary excitatory drive for locomotion by central pattern generators (CPGs) in vertebrates and various invertebrates (Bucher et al., 2015; Kratsios et al., 2024; Von Stetina et al., 2005). For the latter role, primary motor neurons rely on acetylcholine to depolarize target muscle cells through neuromuscular synapses, or junctions (NMJs)(Peper et al., 1982). The NMJs are highly specialized synapses that allow for robust yet exquisite motor control through excitation-contraction coupling. Thus, the gene regulatory networks (GRNs) responsible for the specification and differentiation of cholinergic motor neurons in the central nervous system (CNS) are ones of paramount importance for animal behavior, especially locomotion. Though one might assume these GRNs are largely conserved across metazoans, considerable evidence suggests that invertebrates and vertebrates have evolved varying mechanisms for regulating motor neuron fate and terminal differentiation (Catela et al., 2019; Hobert and Kratsios, 2019; Kratsios et al., 2024; Smith et al., 2024).

Tunicates are non-vertebrate chordates and the sister group to the vertebrates. In the larvae of the tunicate *Ciona robusta*, regulatory mechanisms of cholinergic motor neuron fate specification and differentiation resemble an evolutionary intermediate between vertebrates and other invertebrates. For instance, the transcription factor Ebf acts as a terminal selector for cholinergic fate by activating the expression of a conserved cholinergic locus encoding vesicular acetylcholine transporter and choline acetyl transferase (*VAChT/ChAT*) in Ascending Motor Ganglion neuron 5 (AMG5)(Kratsios et al., 2012; Mathews et al., 2015; Popsuj and Stolfi, 2021), a key interneuron situated just dorsal to the motor neurons of the “core” motor ganglion (cMG, **Figure 1A**)(Kourakis et al., 2025; Ryan et al., 2018). However, it is unclear what regulates cholinergic fate in the motor neurons and other cells of the cMG, as a minimal *VAChT/ChAT* enhancer encompassing a predicted Ebf binding site drives expression in AMG5 but is not required for expression in motor neurons and other cMG cholinergic neurons (Popsuj and Stolfi, 2021). Ebf was recently shown to be necessary for expression of *Nova* in *Ciona* larval motor neurons (Hossain et al., 2025)*. Nova* in turn encodes an alternative splicing factor required for acetylcholine receptor clustering at NMJs in *Ciona* and mammals, through promoting the inclusion of key microexons in *Agrin* (Hossain et al., 2025; Ruggiu et al., 2009).

**Figure 1.**
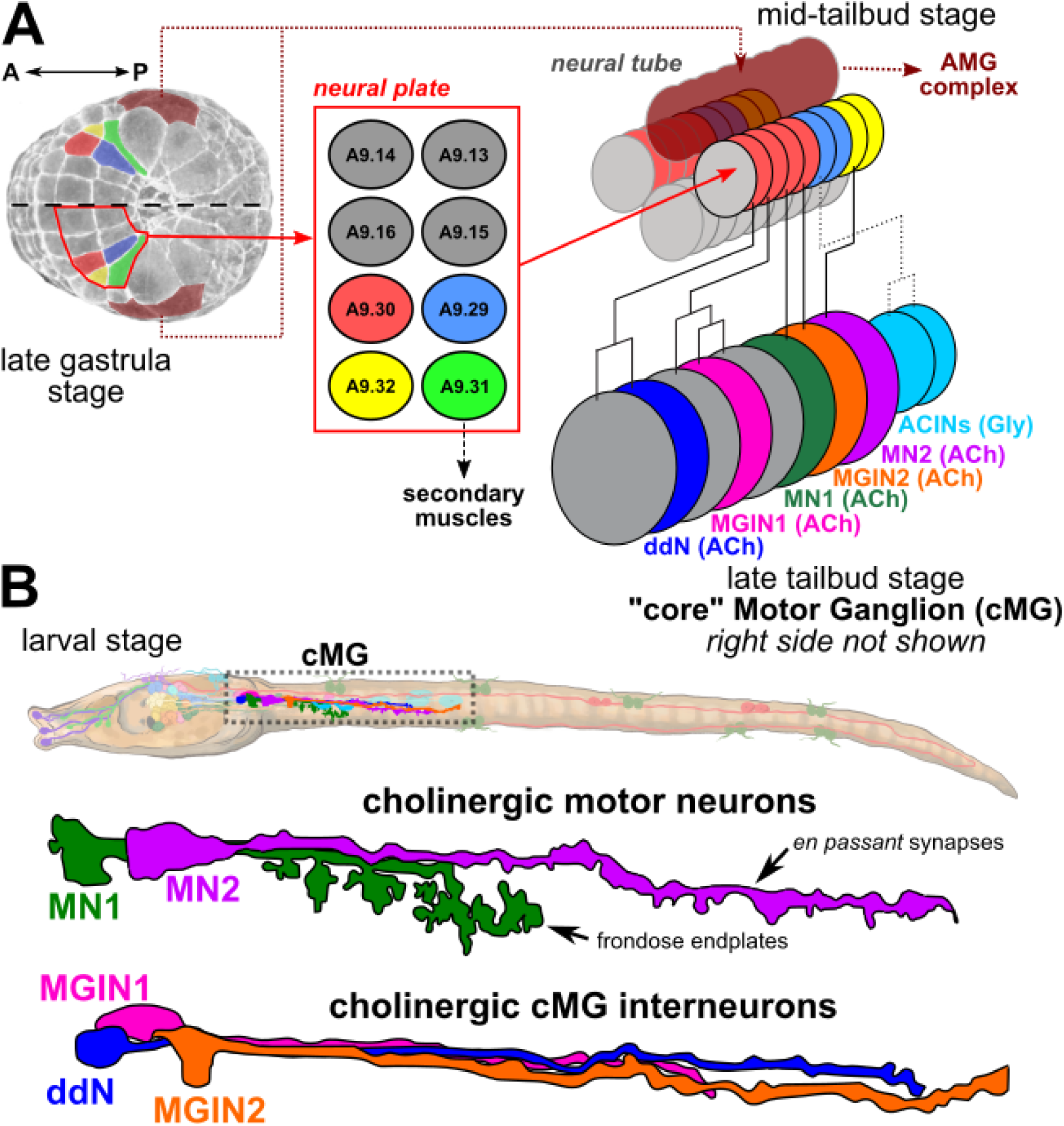
The cholinergic neurons of the “core” motor ganglion of *Ciona robusta*. **A)** Cell lineage diagram depicting the development of the “core” motor ganglion (cMG). AMG: ascending motor ganglion neurons; ACh: cholinergic neuron; ACINs: ascending contralateral inhibitory neurons; ddN: descending decussating neuron; Gly: glycinergic neuron; MGIN: motor ganglion interneuron; MN: motor neuron. **B)** Diagram of larva showing location of cMG neurons. Cholinergic motor neuron and interneuron subtype morphologies shown. Larva illustration by Lindsey Leigh. Neuron morphologies adapted from Ryan and Meinertzhagen (Ryan and Meinertzhagen, 2019).

We sought to understand what is regulating cholinergic fate and other motor neuron differentiation programs in the primary motor neurons and other excitatory cMG neurons in *Ciona*. The cMG, as we define it herein, consists of 5 major left/right bilateral pairs of cholinergic neurons and two bilateral pairs of glycinergic neurons at the base of tail and functions as a CPG for larval motility (Hara et al., 2022). Among the cholinergic neurons, there are two pairs of motor neurons (MN1 and MN2, **Figure 1B**) that form the bulk of the synapses onto the tail muscles of the larva (Ryan et al., 2016). It is thought that these neurons, together with glycinergic inhibitory interneurons located in the anterior part of the tail (the ascending contralateral inhibitory neurons, or ACINs), coordinate most of the tail muscle contractions that drive *Ciona* larval swimming (Hara et al., 2022; Horie et al., 2010; Nishino, 2018; Nishino et al., 2011; Nishino et al., 2010; Zega et al., 2006), although additional motor neurons positioned posteriorly in the tail cord have been recently identified (Kourakis et al., 2025).

Here we report using tissue-specific CRISPR/Cas9-mediated mutagenesis to disrupt different candidate regulatory genes expressed in the cMG. We identified potentially conserved roles for the *Ciona* orthologs of Neurogenin, Ebf, and Onecut transcription factors in promoting differentiation of cholinergic neuron differentiation in the cMG. Moreover, we identified the transcription factor Nkx6 and the secreted Wnt/LRP inhibitor Dkk3 as key regulators of the MN1-specific frondose NMJs, revealing a provisional GRN for motor neuron subtype specification and differentiation in tunicates.

## Results

### Specification of cholinergic neurons in the cMG

To characterize the GRN regulating cholinergic motor neuron fate in *Ciona,* we focused on transcription factors associated with spinal cord motor neuron development in vertebrates and that are also expressed in the developing cMG (Imai et al., 2009; Stolfi and Levine, 2011)(**Figure 2A**). In addition to Ebf, we also focused on the bHLH proneural factor Neurogenin (Neurog), the homeodomain containing transcription factor Onecut, and the Lim homeobox-family factor Lhx3/4. Neurog is upstream of both *Ebf* and *Onecut* in *Ciona*, being necessary for their transcriptional activation in the cMG (Imai et al., 2009). *Ciona* Neurog is the sole *Ciona* ortholog of vertebrate Neurog family members. In vertebrates, Neurog2 induces cholinergic motor neuron fate (Aydin et al., 2019; Hulme et al., 2022; Liu et al., 2013; Mazzoni et al., 2013; Velasco et al., 2017). Similarly, *Ciona* Onecut is orthologous to vertebrate Onecut family members, which are associated with the development of spinal cord neurons, including motor neurons (Audouard et al., 2012; Francius and Clotman, 2010; Roy et al., 2012; Velasco et al., 2017). Finally, Lhx3/4 is orthologous to Lhx3, which is crucial for spinal cord motor neuron development in vertebrates through its activity in a larger transcription factor complex with another LIM homebox factor, Islet1 (Lee et al., 2012; Mazzoni et al., 2013; Seredick et al., 2014). Notably, the sole *Islet* ortholog in *Ciona,* while also downstream of Neurog, appears to only be expressed in one motor neuron subtype, MN2 (Imai et al., 2009; Stolfi and Levine, 2011). Orthologs of all these factors play various roles in neuronal development in different invertebrates, suggesting deeper conservation of GRNs regulating cholinergic and motor neuron fate (Bertrand et al., 2002; Christensen et al., 2020; Leyva-Díaz and Hobert, 2022; Masoudi et al., 2021; Sasakura and Makabe, 2001; Thor et al., 1999).

**Figure 2.**
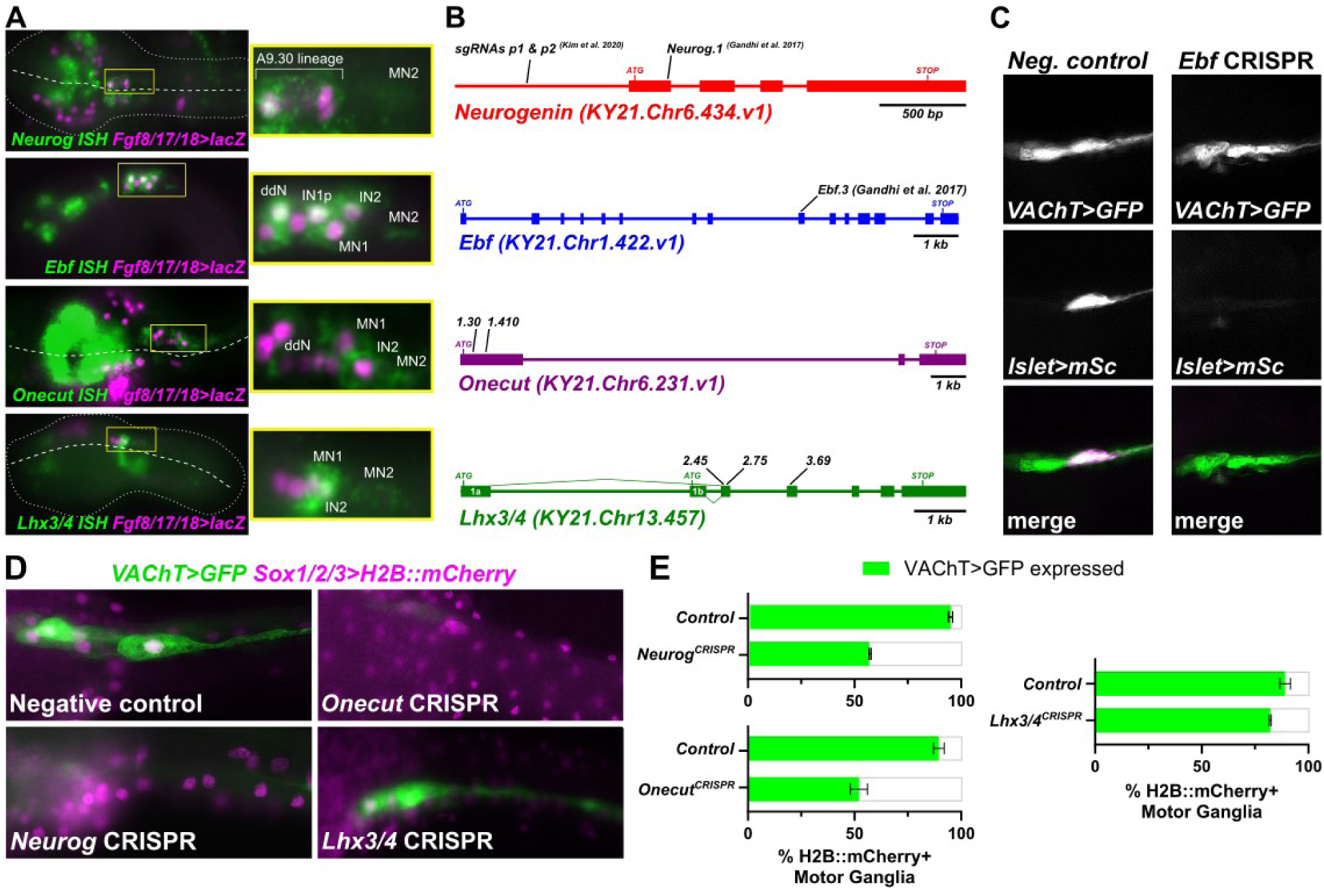
CRISPR/Cas9-mediated mutagenesis of candidate regulatory genes for cholinergic fate. **A)** Fluorescent *in situ* mRNA hybridization (ISH) coupled to immunostaining, showing expression of candidate transcription factor genes in different neuronal subtypes of the cMG. Immunostaining of β-galactosidase driven by the *Fgf8/17/18* promoter reveals the A9.30 cell lineage. ddN: descending decussating neuron; IN1: MG interneuron 1; IN1p: MG interneuron 1 progenitor; IN2: MG interneuron 2; MN1: motor neuron 1; MN2: motor neuron 2. Dotted lines indicate embryo outlines. Dashed line indicated the embryonic midline. **B)** Diagrams depicting candidate regulatory gene loci, indicating sgRNA target sites. Some sgRNAs were previously published. **C)** Neurectoderm-specific CRISPR/Cas9-mediated knockout of *Ebf* does not affect *VAChT -2083/+15>GFP* reporter plasmid expression in the cMG, even though it greatly reduces *Islet* reporter expression (mSc: mScarlet) in MN2, as previously shown (Stolfi et al., 2014). In the control, 48/50 *VAChT>GFP+* larvae also had *Islet>mSc* (magenta) expression in MN2, compared to only 5/50 in the *Ebf* CRISPR condition. **D)** Representative examples of CRISPR/Cas9-mediated mutagenesis of *Neurog, Onecut*, and *Lhx3/4* compared to “negative control” larvae targeted with a non-specific sgRNA. **E)** Graphs depicting scoring of *VAChT -2083/+15>GFP* expression in CRISPR and control larvae represented in panel D. Experiments were performed in duplicate, with n ≥ 22 transfected larvae per condition per duplicate. All error bars show the range between duplicates.

To target these candidate regulatory genes for CRISPR/Cas9-mediated disruption in the cMG, we used the *Sox1/2/3* promoter to drive Cas9 expression in the neurectoderm (Stolfi et al., 2014). We also used previously published and validated single-chain guide RNAs (sgRNAs) targeting *Neurog* (Gandhi et al., 2017; Kim et al., 2020) and *Ebf* (Gandhi et al., 2017), while also designing sgRNAs to target candidates *Onecut* and *Lhx3/4* (**Figure 2B, Supplemental Figure 1**). To assay cholinergic cMG neuron specification and differentiation, we used a GFP reporter plasmid (*VAChT>GFP*), made using a promoter-proximal fragment (-2083/+15)(Hossain et al., 2025) upstream of the conserved cholinergic locus encoding both Vesicular acetylcholine transporter (VAChT) and Choline acetyltransferase (ChAT)(Yoshida et al., 2004). This reporter is active in the cMG and other cholinergic neurons of the central nervous system. Of note, this fragment does not contain the distal AMG5-specific *cis*-regulatory element (*VAChT* - 4315/-3886) bearing a predicted Ebf site and that was previously shown to be activated by Ebf (Kratsios et al., 2012; Popsuj and Stolfi, 2021).

As expected, CRISPR/Cas9-mediated knockout of *Ebf* did not eliminate *VAChT* reporter expression in the cMG, even in larvae that lost MN2-specific *Islet* reporter expression, which has previously been shown to depend on Ebf (Stolfi et al., 2014)(**Figure 2C**). Based on this, we ruled out a role for Ebf in direct transcriptional activation of the minimal *VAChT/ChAT* promoter in the cMG. In contrast, knocking out *Neurog* and *Onecut* both resulted in substantial loss of *VAChT* reporter expression in the cMG, while *Lhx3/4* CRISPR did not have any noticeable effect (**Figure 2D,E**). These data suggested that Neurog and Onecut are necessary for cholinergic fate in the cMG, with Neurog also activating *Onecut* expression in the cMG (Imai et al., 2009). Supporting this, we found that *VAChT/ChAT* reads are substantially upregulated upon Neurog overexpression and downregulated by expression of a dominant-repressor form of Neurog (Neurog::WRPW), by re-analyzing previously published RNAseq data (Kim et al., 2020). Our conclusion from this reanalysis was further supported when we observed dramatic expansion of *VAChT* reporter activity upon overexpression of Neurog (**Supplemental Figure 2**). We thus conclude that, in cMG neurons, Neurog activates Onecut and either or both directly transactivate *VAChT/ChAT*.

### *Cis*-regulatory mutations suggest a major role for Onecut as a direct activator

Given that Neurog is required for Onecut *e*xpression in the cMG (Imai et al., 2009), but both appear to be required for *VAChT/ChAT*, we sought to use *cis*-regulatory mutations to test the relative contributions of either transcription factor. Using the JASPAR database (Rauluseviciute et al., 2023), we found potential Neurog and Onecut binding sites in a smaller *VAChT/ChAT* element (-2083/-742) that was also sufficient to drive reporter expression in the cMG (**Figure 3A,B, Supplemental Sequences File, Supplemental Table 1**). Some of these sites were also found in the corresponding sequence from the related species *Ciona savignyi* (**Supplemental Sequences File**), and no predicted Ebf sites were identified in either species’ sequence, supporting our minimal loss of VAChT/ChAT expression in *Ebf* CRISPR MNs.

**Figure 3.**
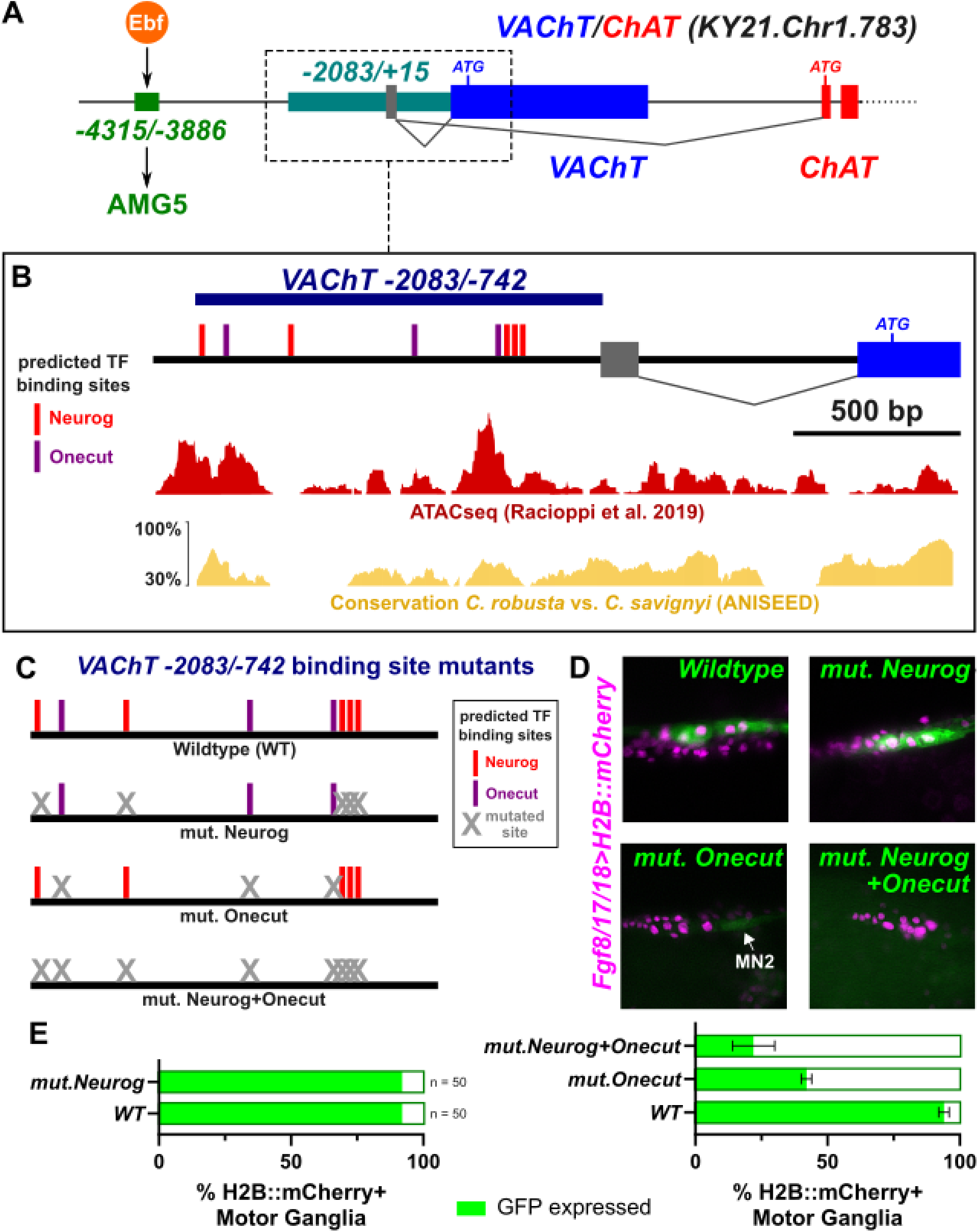
Cis-regulatory analysis of the *VAChT/ChAT* promoter. **A)** Diagram of cholinergic *VAChT/ChAT* locus and different *cis*-regulatory regions active in the AMG5 neuron (Popsuj and Stolfi, 2021), and the remainder of the CNS including cMG cells. **B)** Zoomed in view of the *VAChT -2083/+15* region showing the minimal (to date) fragment driving robust cMG expression (-2083/-742). Candidate transcription factor binding sites predicted by JASPAR are indicated as small colored bars. ATACseq peaks from Racioppi et al. (Racioppi et al., 2019) reproduced from the GHOST genome browser (Satou et al., 2022), and conservation plot from ANISEED (Dardaillon et al., 2020) are further below. **C)** Diagram indicating mutated sites predicted for Onecut or Neurog, within the *VAChT* -2083/-742>GFP reporter, following the color code in panel B. Grey X’s indicate mutated sites. **D)** Representative images of the mutant constructs indicated in panel C, electroporated in larvae together with *Fgf8/17/18>H2B::mCherry* labeling the A9.30 lineage of the cMG (magenta nuclei). No discernible effect on GFP expression (green) was observed upon mutating only predicted Neurog sites. GFP expression was reduced upon mutating predicted Onecut sites, but residual activity was still seen occasionally in MN2 (white arrow). Expression was further reduced by mutating both Neurog and Onecut predicted sites. **E)** Scoring of larvae represented in panel D, as percentage of *Fgf8/17/18>H2B::mCherry*+ cMGs also showing GFP expression. For Onecut and Neurog+Onecut mutant reporters, scoring was performed in biological duplicate and the final percentages averaged, with n = 50 per condition per duplicate. Error bars indicate upper and lower range of percentages between the experimental duplicates.

We tested the *VAChT -2083/-742>GFP* reporter bearing mutations to disrupt predicted Neurog and/or Onecut binding sites (**Figure 3C**). While mutating Neurog sites alone seemed to have little effect on reporter activity, a substantial reduction in GFP signal was observed in all cholinergic neurons of the CNS by mutating predicted Onecut sites (**Figure 3D,E**). However, there was still some reporter activity observed, especially in MN2 (**Figure 3D,E**, **Supplemental Figure 2**). Reporter expression was further reduced in double-mutants in which predicted sites for both Neurog and Onecut were mutated (**Figure 3D,E**). Taken together, these data suggest that Neurog might cooperate with Onecut to ensure robust activation of cholinergic fate in the cMG. Given that Neurog is also necessary for *Onecut* activation in the cMG (Imai et al., 2009) this might constitute a feed-forward loop.

### Nkx6 specifies Motor Neuron 1 (MN1) subtype fate

We next investigated what might be regulating the distinction between the two major motor neuron subtypes in the cMG, MN1 and MN2. These two MN subtypes differ substantially in the size, shape, and positions of the NMJs they form with the tail muscles, all of which are hypothesized to control different aspects of larval motor movements (**Figure 1B**)(Nishino et al., 2011; Ryan et al., 2016). While MN1 forms large, frondose (“leaf-like”) endplates at the base of the tail at its anterior end, contacting the middle band of muscle cells (Imai and Meinertzhagen, 2007; Stolfi and Levine, 2011), MN2 forms *en passant* synapses with the dorsal edge of the muscles, extending further posteriorly into the tail (Hossain et al., 2025; Nishino et al., 2011). Previous work has implicated MN2 in controlling graded muscle activity to produce asymmetric or turning movements (Nishino et al., 2011), and in controlling early tail flicks during hatching (Akahoshi et al., 2021; Utsumi et al., 2023). The role of MN1 is less understood, but this neuron might be involved more heavily in symmetric swimming, based on bilaterally alternating “all-or-none” tail muscle contractions, though MN2 is likely also involved (Akahoshi et al., 2021). Interestingly, MN2 is the only cholinergic neuron of the cMG that does not originate from the A9.30 cell lineage (**Figure 1A**), instead arising from the A9.32 lineage and migrating anteriorly to eventually join the rest of the cMG (Navarrete and Levine, 2016). As such, MN2 maintains some unique differences in its GRN compared to the rest of the cMG. Namely, MN2 uniquely activates motor neuron regulator *Islet* to be expressed only within this A9.32 lineage (Imai et al., 2009; Stolfi and Levine, 2011).

Our primary candidate regulator for MN1 fate was Nkx6 (previously known as Nk6, **Figure 4A**), which was shown previously to be upregulated specifically in MN1, but not MN2, in *Ciona* and other tunicates such as *Molgula* spp (Lowe et al., 2021; Lowe and Stolfi, 2018; Stolfi and Levine, 2011). We also serendipitously identified *Dkk3* (gene ID KY21.Chr8.957) as an MN1-expressed regulatory gene encoding a member of the Dickkopf (Dkk) family of secreted inhibitors of the Wnt signaling pathway (**Figure 4B, Supplemental Figure 3**). To visualize MN1 neurons, we used reporter plasmids (*Claudin.j>GFP or mScarlet*) constructed using the first intron of *Claudin.j* (previously named *Claudin7,* gene ID KY21.Chr10.463), which drives robust fluorescent reporter gene expression in MN1 and its endplates, consistent with the strong *Claudin.j in situ* mRNA hybridization signal in this cell (**Figure 4C,D**).

**Figure 4.**
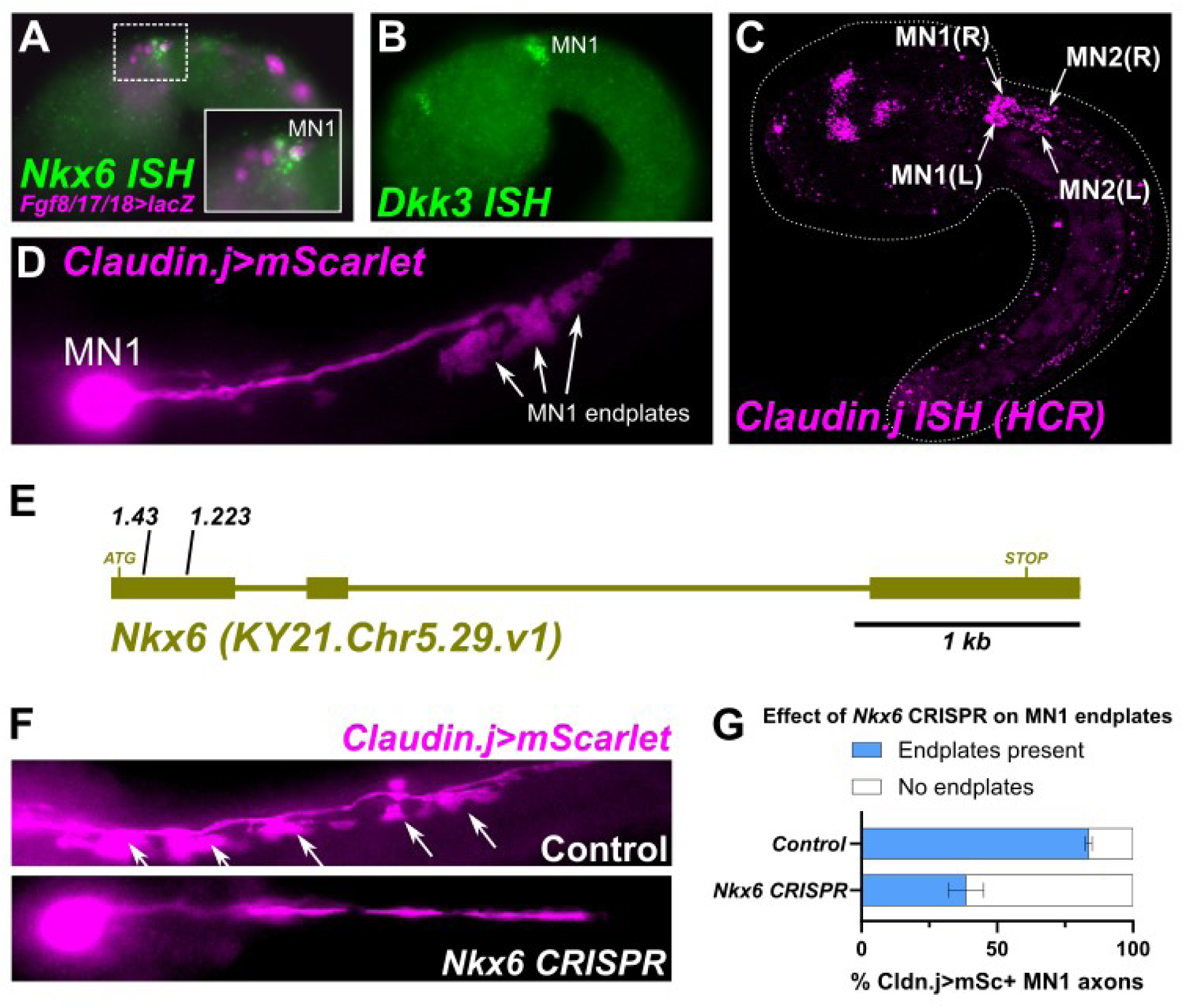
Regulation of MN1 differentiation by Nkx6. **A)** Whole-mount fluorescent mRNA *in situ* hybridization (ISH) and immunostaining showing expression of *Nkx6* in MN1. **B)** *In situ* hybridization of *Dkk3* showing expression in MN1. **C)** Hybridization chain reaction (HCR) *in situ* showing expression of *Claudin.j* mRNAs in diverse neurons including motor neurons. Expression is observed in both MN1 and MN2, but highly enriched in MN1. Dorsolateral view showing both left (L) and right (R) sides. **D)** Fluorescent reporter constructed using the first intron of *Claudin.j* reveals the frondose endplates typical of MN1. **E)** Diagram of *Nkx6* locus, showing target sites of *Nkx6* sgRNAs. **F)** Representative images showing effect of *Nkx6* CRISPR on MN1 endplate formation, showing absent or highly reduced endplate formation in the CRISPR condition (*Fgf8/17/18* promoter was used to drive Cas9 expression specifically in the A9.30 cell lineage). *Claudin.j* reporter is unaffected by *Nkx6* CRISPR, allowed for visualization of the cell body and axon. **G)** Graph of scoring *Nkx6* CRISPR and negative control larvae for presence or absence of MN1-type endpates. Experiment performed in duplicate, n ≥ 40 neurons per condition, per duplicate. Error bars show range between duplicates.

To test the role of Nkx6 in regulating MN1 development, we designed and validated new sgRNAs targeting the gene (**Figure 4E, Supplemental Figure 1**). We selected the *FGF*8/17/18 promoter to confine our CRISPR/Cas9-mediated disruption to cells within the A9.30 lineage (which gives rise to MN1)(Imai et al., 2009). Upon *Nkx6* CRISPR, *Claudin.j>mScarlet* expression was not lost, allowing us to visualize MN1 cell bodies and motor endplates. Strikingly, we observed a high frequency of larvae that no longer had the distinctive frondose endplates of MN1 present (**Figure 4F,G**). While the cell bodies and axons of MN1 were still visible as labeled by *Claudin.j>mScarlet*, the MN1 endplates were absent or greatly reduced (**Figure 4F,G**). These results suggest that Nkx6 is required for the development of MN1-specific motor endplates in the *Ciona* larva.

### CRISPR-mediated disruption of *Dkk3* also abolishes MN1 endplates

Given that Nkx6 is a transcription factor, it is unlikely it is acting directly as a molecular mechanism for endplate formation. Rather, we hypothesized that one or more effectors might also be involved. In support of this hypothesis, we identified Dkk3 as a unique MN1-specific marker potentially involved in endplate formation. Although its very early initial expression in MN1 suggested Dkk3 is unlikely to be downstream of Nkx6, we reasoned it was an intriguing candidate effector based on its membership in the Dkk family of Wnt pathway inhibitors (Cruciat and Niehrs, 2013; Niehrs, 2006). The Dkk family overall is best known for its ability to competitively inhibit the Wnt-Frizzled complex by binding to the LRP co-receptors thereby preventing Frizzled/LRP interaction induced by Wnt ligand binding. (Bao et al., 2012; Brott and Sokol, 2002; Niehrs, 2006; Wen et al., 2023). LRP receptors additionally play conserved roles in NMJ formation (DePew et al., 2024; Weatherbee et al., 2006; Zong et al., 2012), including in the *Ciona* larva, in which Lrp4 was shown to be required for acetylcholine receptor clustering at NMJs (Hossain et al., 2025). Additionally, more recent studies posit a role for Wnts and their inhibitors in impacting NMJ formation, leading us to hypothesize a possible role for Dkk3 within MN1 (Boëx et al., 2018; DePew and Mosca, 2021; Shen et al., 2018).However, the role of the Dkk family in regulating NMJ formation is still unclear.

We aligned the *C. robusta* Dkk3 protein sequence to Dkk3 sequences from various chordate species (**Figure 5**). We found that *Ciona* and all vertebrate Dkk3 orthologs possess at least one, sometimes multiple NxI/V motifs (where N = asparagine, “x” represents any amino acid, and I/V = isoleucine or valine) that are key for Dkk1 and Agrin ligands to bind to LRP receptors (Zong et al., 2012). We also found that *Ciona* Dkk3 had two additional NxF motifs (F = phenylalanine), which can substitute for NxI/V in *C. robusta* Agrin (Hossain et al., 2025). We therefore selected Dkk3 for further investigation as a candidate effector of MN1 endplate development in *Ciona*.

**Figure 5.**
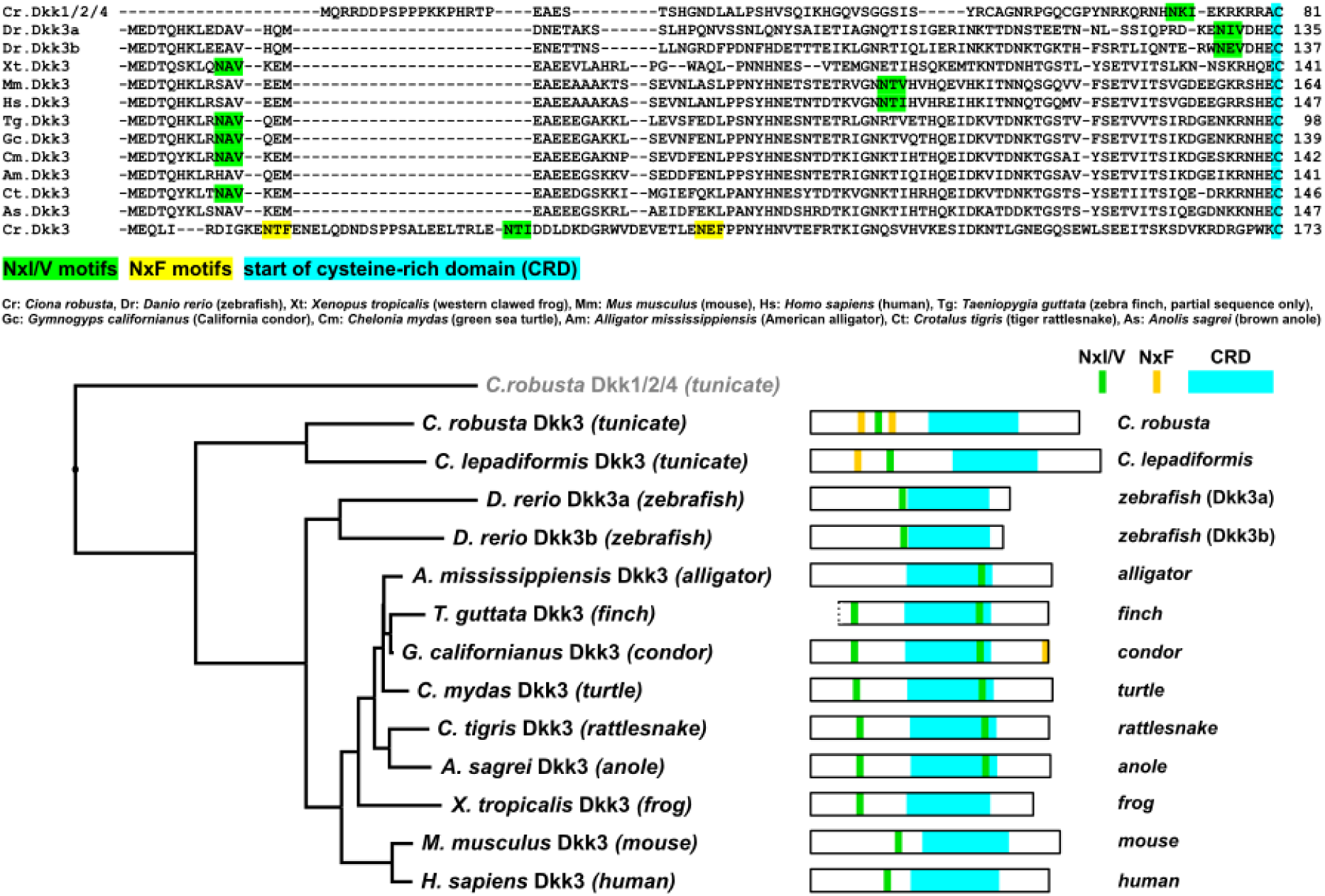
Dkk3 orthologs in the chordates. Top: alignment of Dkk3 orthologs from various chordate species, as seen below in Dkk3 phylogenetic tree. NxI/V motifs, NxF motifs, and cysteine-rich domains (CRDs) color-coded according to the legend. Bottom: phylogenetic tree of Dkk3 orthologs, and diagrams depicting motif and domain organizations.

To knock out *Dkk3*, we used a previously validated sgRNA targeting the gene (Popsuj et al., 2024), and selected the *Sox1/2/3* promoter to drive expression of Cas9 in neural progenitors. We used the *Claudin.j>mScarlet* reporter to visualize MN1 endplates, and *Tbx6-r.b>AchRA1::GFP* to visualize acetylcholine receptor clusters in the muscles (Hossain et al., 2025; Nishino et al., 2011). Like for the *Nkx6* CRISPR, we observed a loss or severe reduction of MN1 endplate formation upon *Dkk3* CRISPR (**Figure 6A,B**). We found a significant reduction in the average number of endplates per MN1 axon in *Dkk3* CRISPR larvae in three out of four biological replicates, largely due to the number of larvae in which MN1 endplates were completely absent (**Figure 6C**). Although we found no evidence to suggest that Dkk3 impacts endplate size (**Figure 6D**), we observed a significant reduction in AChR clustering at MN1 endplates in all four biological replicates (**Figure 6F,G**). Notably, AChR clusters could still be seen at MN2 synapses, suggesting a specific effect in MN1, where Dkk3 is expressed (**Figure 6G**). These results indicate that *Dkk3* is required for the formation of normal synapses between MN1 and its target muscle cells.

**Figure 6.**
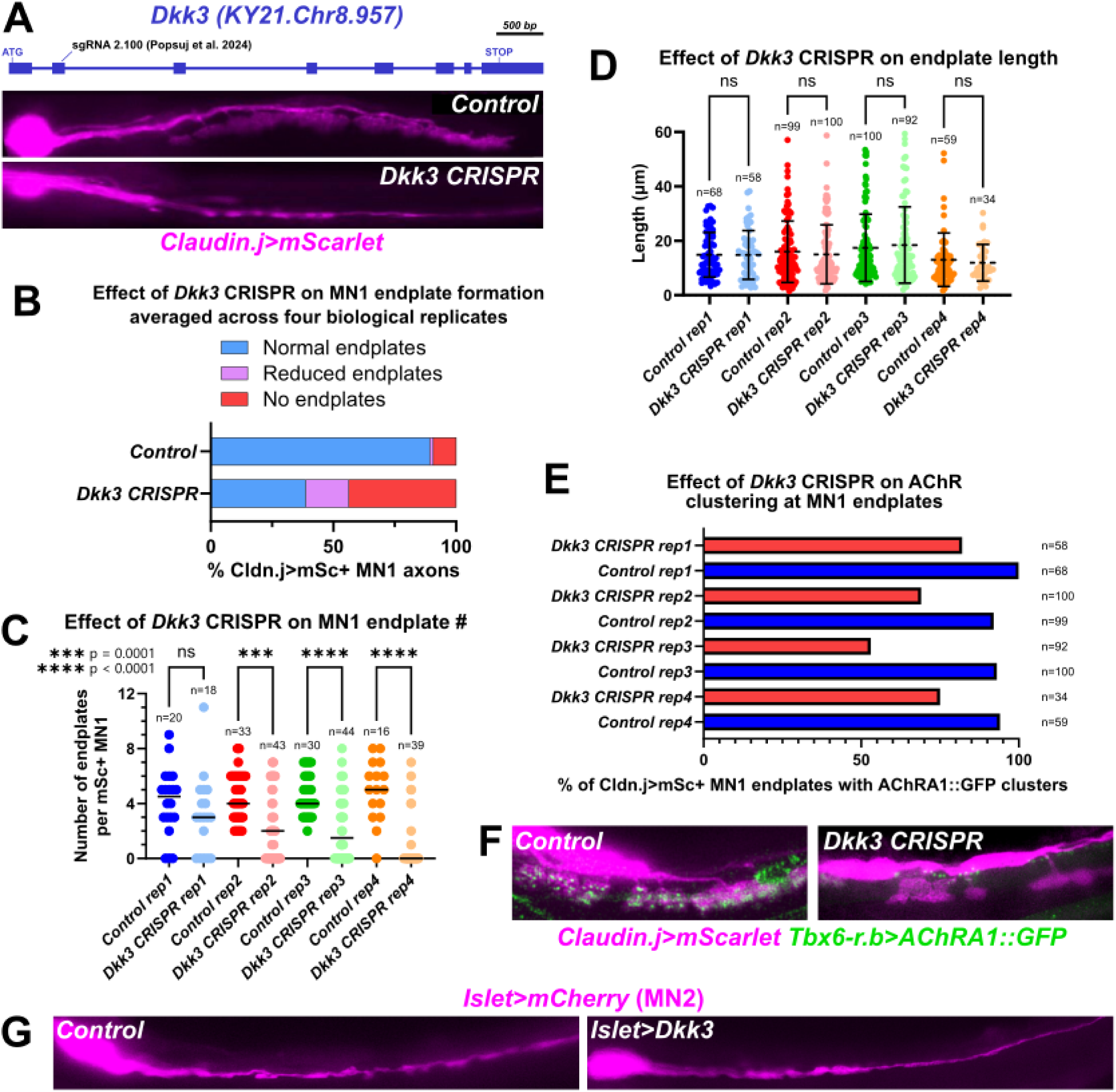
Regulation of MN1-type endplate formation by Dkk3. **A)** *Dkk3* locus diagram showing published sgRNA target. Bottom: representative images showing loss/reduction of MN1 endplates upon *Dkk3* CRISPR. **B)** Graph showing effect of *Dkk3* CRISPR on MN1 endplate formation, from data across four independent replicates, n ≥ 15 larvae per condition, per replicate. The *Sox1/2/3* promoter was used to drive Cas9 expression in the neurectoderm. **C)** Plot showing effect of *Dkk3* CRISPR on the number of endplates observed per visible MN1 axon. Statistical tests calculated as two-tailed Mann-Whitney tests, ns = not significant. **D)** *Dkk3* CRISPR did not have a significant effect on the length of those MN1 endplates that remained. Statistical test calculated as two-tailed Mann-Whitney test. **E)** Graph showing scoring of acetylcholine receptor (AChRA1::GFP) clusters in the tail muscles in contact with MN1 endplates, in *Dkk3* and control larvae. **F)** Representative images of presence or absence of AChRA1::GFP clusters (green dots) in *Dkk3* CRISPR or control larvae, as plotted in panel E. AChRA1::GFP clusters in the *Dkk3* CRISPR condition can still be seen at MN2 synapses, along the dorsal edge of the muscles. **G)** Representative images of MN2 labeled with *Islet>mCherry* (magenta), showing no discernable presence of MN1-type endplates upon mis-expression of Dkk3 in this neuron. Experiment was replicated thrice. Quantification in panels B-E obtained from same set of images.

We next asked if Dkk3 is sufficient to induce MN1-type frondose endplates when mis-expressed in MN2. To do this, we used a *cis-regulatory* element from *Islet* (Stolfi et al., 2010) that drives expression in a subset of CNS neurons including MN2 (*Islet[CNS]>Dkk3)*. However, we did not observe any ectopic, MN1-like frondose endplates in MN2 axons upon electroporation with *Islet[CNS]>Dkk3*, even across three biological replicates (**Figure 6H**). This suggests that Dkk3 is not sufficient for establishing MN1-type frondose endplates, indicating that there are other required MN1-specific effectors yet to be identified, or that there are unknown MN2-specific factors inhibiting frondose endplate formation in these neurons.

### *Ciona* Dkk3 can alter Wnt/LRP signaling through NxI/V/F motifs

To better understand the mechanism by which Dkk3 could be promoting endplate formation in MN1, we next investigated its potential modulation of Wnt/LRP activity. Given the ability of Dkk family members to bind LRP receptors, and the presence of three NxI/V/F motifs in *Ciona robusta* Dkk3, we hypothesized that it could be binding to Lrp4 in tail muscle cells via these motifs, in a manner similar to motor neuron-derived Agrin binding to Lrp4 via its NxF motif to promote acetylcholine receptor clustering, as recently suggested in *Ciona* (Hossain et al., 2025). However, in the absence of a gain-of-function phenotype in motor neurons upon Dkk3 overexpression (**Figure 6G**), we were unable to easily test this in the context of NMJ development. With this in mind, we devised a binary assay to assess the competence of Dkk3 to bind LRP receptors in the context of competitive inhibition of Wnt/LRP signaling *in vivo* (**Figure 7A**). The two pigment cells of the *Ciona* larvae (ocellus and otolith) arise from a bilaterally symmetric equivalence group. During neural tube closure, these cells undergo random intercalation at the dorsal midline; the cell that occupies the more posterior position contacts the posterior nerve cord receives a Wnt7 signal (“Wnt on”) and become the ocellus. Conversely, the cell that ends up more anterior fails to receive this signal (“Wnt off”), and instead differentiates as the otolith cell (Abitua et al., 2012).

**Figure 7.**
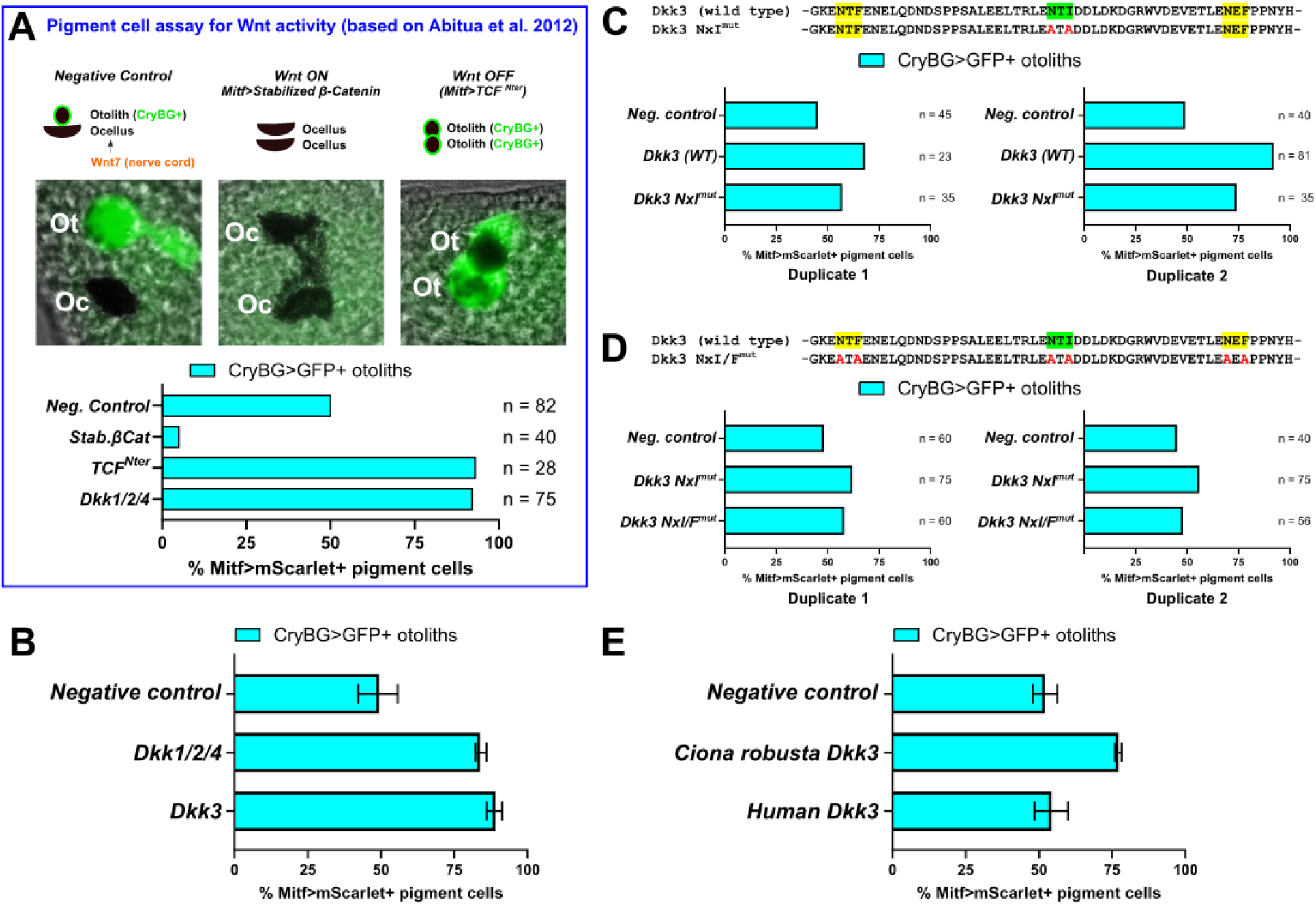
A pigment cell assay reveals inhibition of LRP/Wnt signaling by Dkk3. **A)** Expected outcomes and results of a proof-of-principle experiment establishing an otolith (Ot) and ocellus (Oc) pigment cell-based assay for LRP/Wnt inhibition, based on findings from Abitua et al. (Abitua et al., 2012). See text for details. **B)** Pigment cell assay reveals comparable inhibitory effect of *Ciona robusta* Dkk3 and Dkk1/2/4. Experiment performed in triplicate, n ≥ 65 pigment cells per condition, per replicate. **C)** Mutating the NxI motif in *Ciona* Dkk3 lowers its inhibitory activity compared to the wild type (WT) Dkk3, as assayed in duplicate. **D)** Mutating the two NxF motifs further reduces the inhibitory effect of *Ciona* Dkk3, as assayed in duplicate. **E)** Human Dkk3 does not have the same ability as *Ciona robusta* Dkk3 to inhibit LRP/Wnt signaling in our pigment cell assay. Experiment performed in duplicate, n ≥ 16 in control conditions, n ≥ 35 in Dkk3 conditions. All error bars indicate range across duplicates/replicates.

As a proof-of-principle, we used the *Mitf* promoter (Abitua et al., 2012) to drive pigment cell-specific expression of either stabilized β-catenin (Wada et al., 2008), which should hyperactivate canonical Wnt signaling, or the N-terminus of TCF (TCF^Nter^), which acts as an inhibitor of the pathway (Kaplan et al., 2019). To monitor otolith specification, we used the *CryBG>H2B::GFP* reporter (Shimeld et al., 2005), which is expressed in the otolith pigment cell but not the ocellus pigment cell. In the negative control population (no Wnt manipulation), roughly 50% of Mitf+ pigment cells are fated to become otoliths and express the CryBG reporter (**Figure 7A**). Therefore, CryBG+ otolith frequency over ∼50% across the population indicates a repressive effect on Wnt signaling (more otoliths, fewer ocelli), while frequencies under ∼50% indicate hyperactivation of Wnt (fewer otoliths, more ocelli). Indeed, TCF^Nter^ the conserved Wnt/LRP inhibitor, Dkk1/2/4, showed a high degree on Wnt-repressive activity in this assay, while stabilized β-catenin showed a clear hyperactive Wnt phenotype, therefore confirming the validity of our assay (**Figure 7A**).

After establishing this assay, we next sought to test the effect of overexpressing different Dkk3 variants on Wnt/LRP signaling in the pigment cell lineage. First, we found that wild-type *Ciona* Dkk3 can indeed inhibit Wnt signaling in this context, confirming its ability to interfere with the pathway in a manner roughly comparable to Dkk1/2/4 (**Figure 7B**). To test the hypothesized role the NxI motif present in *Ciona* Dkk3 we mutated it to AxA instead (A = alanine). Although this mutation reduced the Wnt-repressive activity of Dkk3, it did not completely abolish the ability to inhibit Wnt signaling (**Figure 7C**).

However, as we mentioned above, there are also two NxF motifs in *Ciona* Dkk3 (**Figure 5**) which we hypothesized could substitute for NxI/V as in *Ciona* Agrin. We therefore mutated all three NxI/F motifs in *Ciona* Dkk3 and tested its activity in our pigment cell assay. Indeed, we measured a further reduction of Wnt inhibition of this triple NxI/F mutant variant compared to the single NxI mutant (**Figure 7D**). However, otolith frequency was not quite at 50%, suggesting additional domains through which Dkk3 could still exert residual inhibitory activity on Wnt/LRP signaling, like for instance the conserved cysteine-rich domain, which has also been implicated in LRP binding and modulation of Wnt pathway activation (Brott and Sokol, 2002; Li et al., 2002).

Given contradictory results in the literature regarding the competence of vertebrate Dkk3 to bind LRP and/or inhibit Wnt signaling (Mourtada et al., 2023), we also tested the human Dkk3 protein sequence in this pigment cell assay. However, did not find any significant effect of human Dkk3 on Wnt/LRP signaling when tested side-by-side with *Ciona* Dkk3 in our assay (**Figure 7E**). This suggests that human Dkk3 might not interact with LRP receptors in the same way, or that there has been considerable co-evolution between Dkk3 and LRP receptors in different chordate groups. Taken together, our data suggest that, indeed *Ciona* Dkk3 can modulate Wnt/LRP activity, likely through its three N-terminal NxI/V/F motifs, but this mechanism might not be conserved in mammals.

## Discussion

We have presented emerging evidence for a GRN regulating cholinergic neuron specification and differentiation in the cMG of the *Ciona* larva (**Figure 8**). While Ebf appears to act as a terminal selector of cholinergic fate in AMG5, a cholinergic interneuron in the dorsal Ascending Motor Ganglion (AMG) complex, it likely does not play the same role in the cMG. Instead, our data point to a major role for Neurog and Onecut in activating *VAChT/ChAT* transcription. However, Ebf has been shown to play an indispensable role in directly activating the transcription of *Nova* in *Ciona* motor neurons, which is required for the conserved, but chordate-specific alternative splicing event that allows for motor neuron-derived Agrin to induce acetylcholine receptor clustering at the NMJs (Hossain et al., 2025). Interestingly, disrupting predicted Onecut-binding sites in the *VAChT/ChAT* promoter appeared to have a greater deleterious effect in A9.30 lineage-derived neurons than in MN2, which is derived from the neuromesodermal progenitors of the A9.32 lineage instead (Navarrete and Levine, 2016). This potentially points to slightly different regulatory logic operating in the two lineages, with Onecut being more crucial for cholinergic expression in the A9.30 lineage than in MN2. Evidence has recently been presented suggesting most of the cMG is homologous not to the vertebrate spinal cord, but rather to the hindbrain (Kourakis et al., 2025). The distinct cell lineage history and regulatory logic of MN2 suggests this neuron subtype might be homologous to vertebrate spinal cord neurons, while MN1 and the rest of the cholinergic cMG neurons are indeed homologous to hindbrain neurons instead.

**Figure 8.**
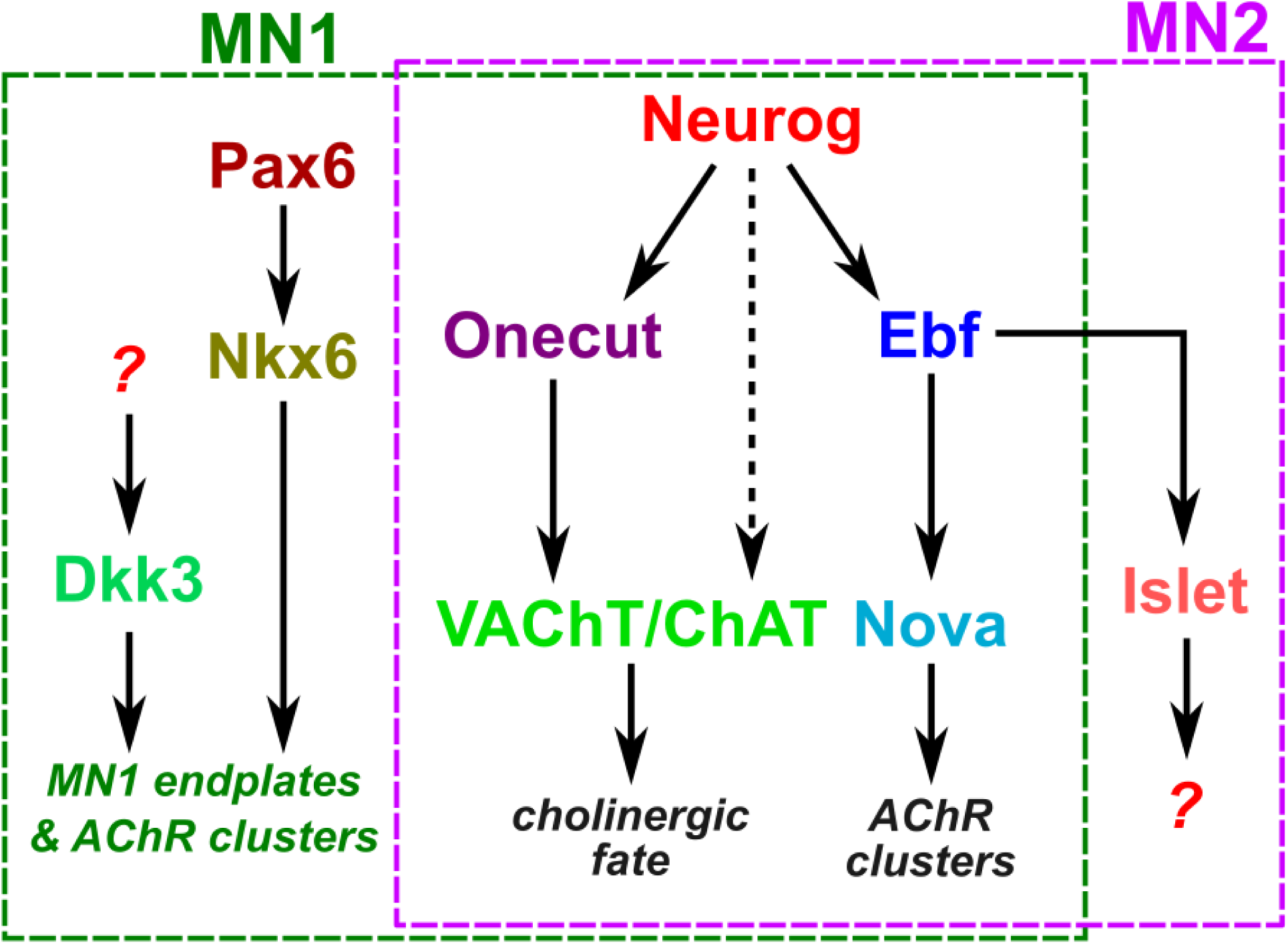
Proposed gene regulatory network for motor neuron specification and differentiation in the *Ciona* larva. Dotted lines indicate a potential direct role for Neurog in a feed-forward loop for sustained expression of *VAChT/ChAT* and/or an indirect role through Onecut, depending on the cMG neuron subtype (see text for details). Question marks indicate unknown targets and function (Islet) or regulators (of Dkk3). Connections between Pax6-Nkx6, Neurog-Onecut and Neurog-Ebf are from Imai et al. 2009. Connection between Ebf-Islet is from Stolfi et al. 2014. Connection between Ebf-Nova is from Hossain et al. 2025.

This regulatory logic downstream of Neurog is somewhat consistent with what is known in vertebrates, in which Neurog2 is necessary to begin chromatin-remodeling steps necessary for transcription of motor neuron-associated genes, also through its activation of Ebf and Onecut factors (Francius and Clotman, 2014; Roy et al., 2012; Velasco et al., 2017). In the case of vertebrate motor neurons, the activation of Ebf and Onecut through Neurog is also essential for key motor neuron transcription factors Islet and Lhx3 to bind to downstream enhancers. We did not find any evidence for a substantial contribution of Lhx3/4 to cholinergic fate in *Ciona* larval motor neurons. It is worth noting that, in the *Ciona* cMG, Islet is an MN2-specific marker but does not appear to be meaningfully expressed in MN1 or other cholinergic MG neurons. However, the low level of GFP expression seen with the VAChT/ChAT reporter plasmid lacking both Neurog and Onecut sites suggests additional factors, like Lhx3/4 or Islet, could be playing a supportive role. It is also possible that vertebrate-specific changes could have contributed to a greater variety of motor-neuron subtypes present in vertebrates, requiring a more complex GRN with additional regulatory inputs. In contrast, the *Ciona* cMG cholinergic neuron GRN might be a simplified one associated with the rapid specification and differentiation of larval motor neurons in the short-lived larval phase of its life cycle. It remains to be seen whether this larval GRN is redeployed as is, or further modified, during the specification and differentiation of adult motor neurons.

We have also revealed in this study a new regulatory pathway controlling the development of MN1-specific neuromuscular synapses. While MN1 and MN2 still share a common gene regulatory network for establishing cholinergic identity and acetylcholine receptor clustering, but differ in the size, shape, and position of their NMJs. We have now identified the MN1-specific upregulation of the transcription factor Nkx6 as likely a key step towards forming the unique, frondose endplates that are characteristic of this motor neuron subtype in the tunicate larva. We believe this will be of great interest to researchers working on MG neuron function, since the exact role of MN1 in swimming behavior has yet to be definitively shown, and its functions are not as well understood as those of MN2 (Akahoshi et al., 2021; Nishino et al., 2011).

We have further identified a novel role for Dkk3 as a regulator of MN1-specific endplates, which we propose is due to its ability to bind LRP receptors in target muscle cells. The onset of *Dkk3* expression in MN1 appears to be concurrent with or earlier than that of *Nkx6,* which suggests parallel mechanisms in play. Although this is the first report, to our knowledge, of Dkk3 impacting any aspect of neuromuscular synapse or endplate formation, it has been shown that Dkk1 can bind to LRP receptors in cultured mouse myotubes (Koles and Budnik, 2012; Messéant et al., 2017; Wang et al., 2008). Although some results suggested that Dkk1 inhibits acetylcholine receptor clustering in cultured myotubes (Messéant et al., 2017), others suggest that it promotes clustering through inhibition of canonical Wnt signaling (Wang et al., 2008). These discrepancies might be due to the complex roles of canonical and non-canonical Wnt in vertebrate NMJ development (Koles and Budnik, 2012).

Although we identified a key role for N-terminal NxI/V/F motifs in the ability of Dkk3 to modulate the Wnt and/or LRP receptors, we did not find evidence that this feature is conserved in mammalian Dkk3. In Dkk3 protein sequence alignments, human and mouse Dkk3 have a single NxI motif in the N-terminal region, however it is not in a conserved location relative to other vertebrate Dkk3 proteins. It is possible that mammalian Dkk3 has lost the ability to strongly bind to LRP and inhibit Wnt signaling through competitive binding. Alternatively, there may have been substantial co-evolution between Dkk3 and LRP receptors in tunicates and/or vertebrates. Of note, the neuron-specific “Z” exons in *Ciona robusta Agrin* encode an NxF motif in place of the NxI motif encoded human and mouse *Agrin* Z exons (Hossain et al., 2025). This may reflect a selective advantage for specifically NxF motifs (not NxI/V) in *C. robusta.* Future studies with a wider range of vertebrate Dkk3 orthologs will be needed to examine the possibility of NxI/V/F motif-receptor coevolution. In *Ciona, Dkk3* is expressed in diverse neuronal types and even the notochord (**Supplemental Figure 3**), suggesting a diversity of functions beyond neuromuscular synapse development. The variable number, position, and exact composition of NxI/V/F motifs found in Dkk3 orthologs in various chordate groups suggests dynamic evolution of the Dkk3 gene, which has potentially been co-opted to regulate diverse Wnt/LRP-dependent processes in different species.

## Materials and methods

### General embryo handling and electroporation

*Ciona robusta* (also known as *Ciona intestinalis* Type A) adults were collected and shipped by M-REP (Carlsbad, CA) or Marinus Scientific (Lakewood, CA). Adult specimens were dissected to obtain gametes for *in vitro* fertilization and subsequent dechorionation and electroporation as previously described (Christiaen et al., 2009a, b). When electroporating reporter plasmids, 15-90 µg of plasmid DNA per 700 µl of total electroporation volume was used depending on promoter strength and specific localization tag (e.g. H2B, Unc-76, etc.) used. For Cas9/Cas9::GemN, sgRNA, and over/mis-expression constructs, 40-100 µg was electroporated instead. Relevant sgRNA, *cis-*regulatory, and protein-coding sequences can be found in the **Supplemental Sequences** file. *VAChT* promoter sequences were analyzed for candidate transcription factor binding sites on JASPAR (https://jaspar.elixir.no/) following the default settings (Rauluseviciute et al., 2023). Larvae were reared to 17 hours post-fertilization (hpf) at 20°C (Hotta Stage 27)(Hotta et al., 2020) in all assays unless otherwise specified.

### Design and validation of sgRNAs for CRISPR

CRISPR sgRNA designs were performed using CRISPOR (https://crispor.gi.ucsc.edu/) (Concordet and Haeussler, 2018; Haeussler et al., 2016), following *Ciona*-specific protocols and guidelines previously (Popsuj et al., 2024). Validation of unpublished sgRNAs were carried out as described (Popsuj et al., 2024), using the Amplicon-EZ service by Azenta. Tissue-specific CRISPR knockouts in F0 were carried out using various promoters to drive restricted expression of Cas9 or Cas9::Geminin^N-terminus^ (Cas9::GemN) as described (Popsuj et al., 2024). All sgRNA targeting sequences and primers used for amplicon PCR for validation shown in the **Supplemental Sequences** file.

### Fixing, mounting, imaging

Embryos and larvae were fixed in MEM-formaldehyde solution and mounted in 1X PBS, 2% DABCO, 50% glycerol mounting solution as previously described (Popsuj et al., 2023). All images were acquired using a Leica DMI8 or DMIL LED inverted compound epifluorescence microscope, or a Zeiss LSM 900 laser scanning confocal microscope. Post-imaging analysis of images acquired on Leica microscopes was performed using the Leica LASX software, including measurements of MN1 endplate lengths. Images obtained on the Zeiss confocal were processed and analyzed using ZEN software.

### mRNA *in situ* hybridization

*Neurog, Ebf, Onecut, Lhx3/4, Ebf, Nkx6,* and *Dkk3 in situs* were performed as previously described (Beh et al., 2007; Ikuta and Saiga, 2007; Stolfi et al., 2011). Probe template sequences or gene collection IDs (Satou et al., 2002) are included in the **Supplemental Sequences** file. For *Claudin.j,* hybridization chain reaction (HCR) was performed following the protocol recommended for marine invertebrates (Bruce et al., 2021). Proprietary probes and hairpins for B1 amplification were generated by Molecular Instruments.

## Supporting information

Supplmental Sequences

## Acknowledgments

We would like to acknowledge Susanne Gibboney, Lindsey Cohen, and Joshua Kavaler for technical assistance. This research was supported by NIH grants R35GM158421 to AS and R01HD104825 to CDC and AS, and by fellowships from ARCS and Mortar Board to SP.

---

**Supplemental Figure 1.**
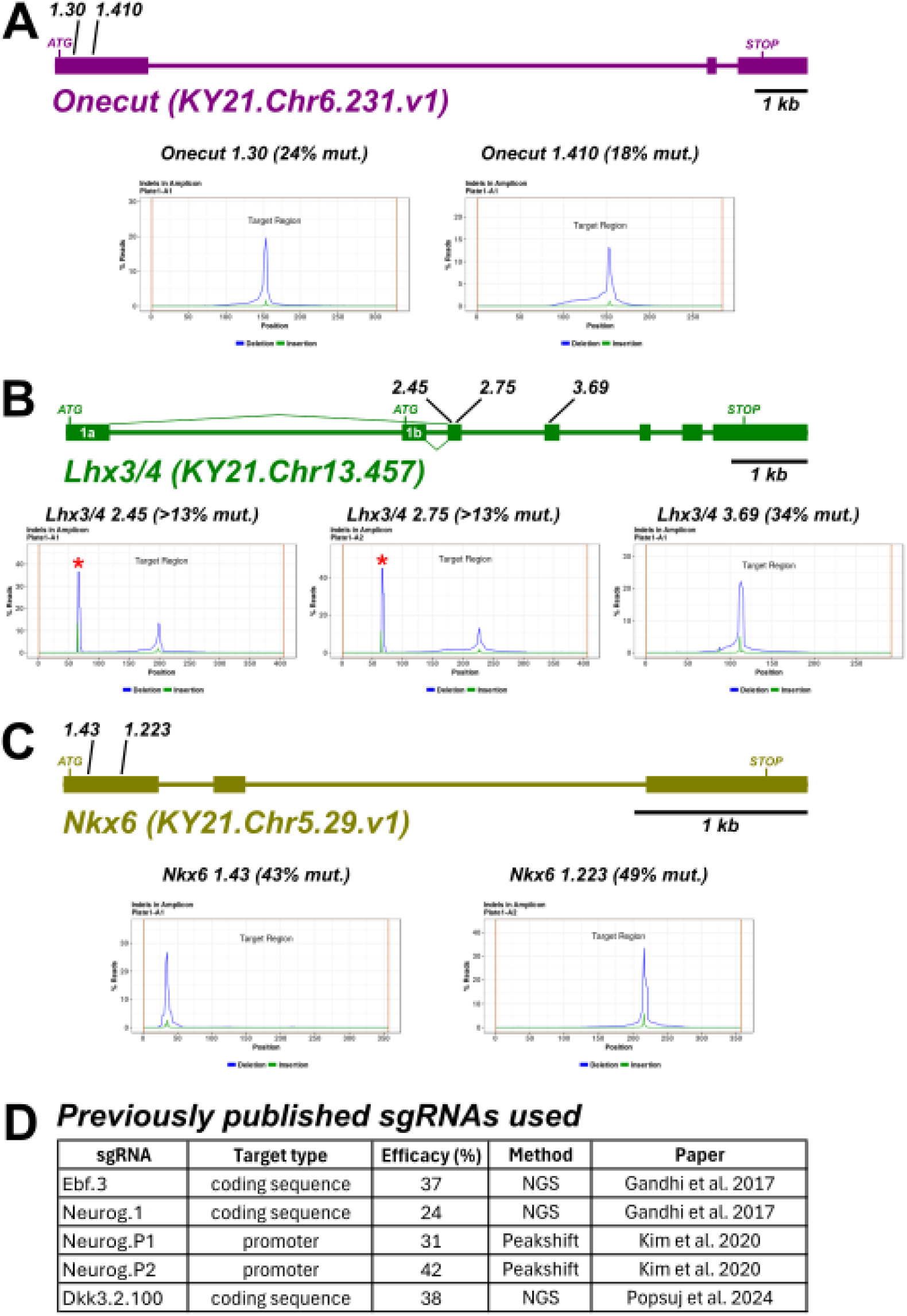
Validation of sgRNAs used. Validation of novel sgRNAs targeting **A)** *Onecut*, **B)** *Lhx3/4*, and **C)** *Nkx6*, using Illumina sequencing of target site amplicons. Asterisks denote naturally occurring indel. **D)** Summary of validation efforts for published sgRNAs used in this study.

**Supplemental Figure 2.**
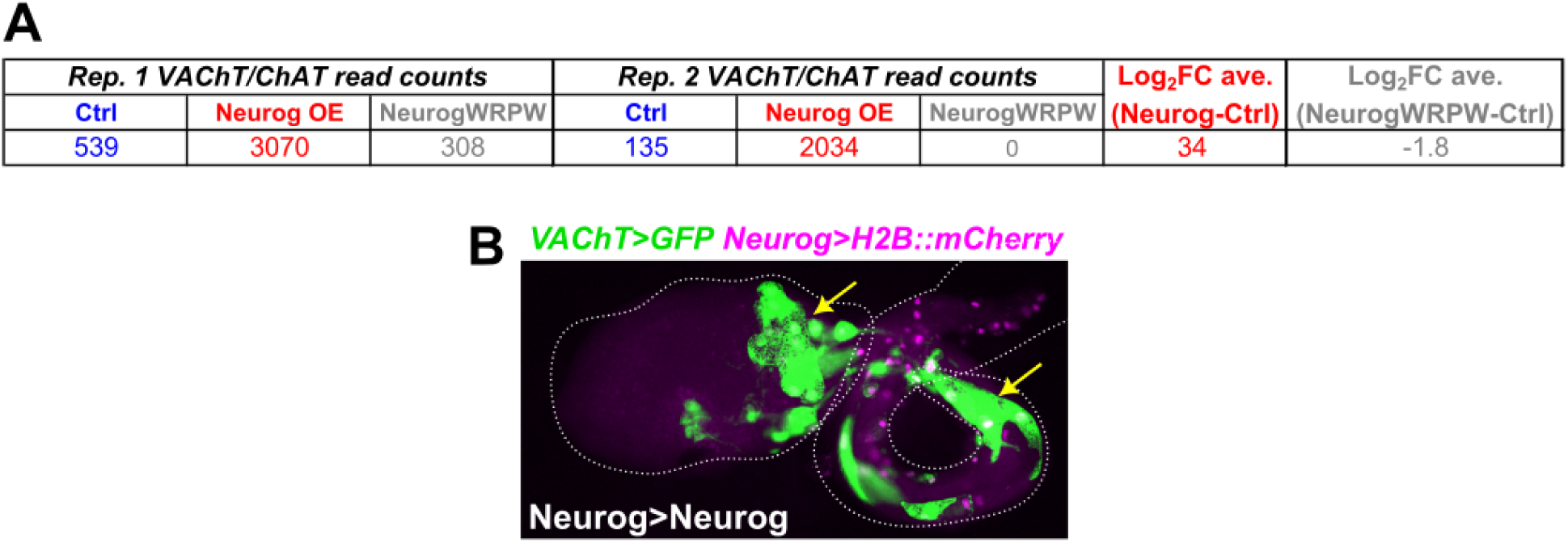
RNAseq analysis of effect of Neurog manipulations on *VAChT/ChAT* expression. **A)** Table showing RNAseq data from Kim et al. 2020, comparing number of *VAChT/ChAT* reads between negative control (“Ctrl”), Neurog overexpression (Neurog>Neurog, “Neurog OE”), and Neurog::WRPW expression (Neurog>NeurogWRPW), all in FACS-isolated bipolar tail lineage cells. *VAChT/ChAT* is upregulated by Neurog overexpression, and downregulated by expressing the dominant repressor NeurogWRPW. **B)** Overexpression of Neurog the bipolar tail neuron lineage-specific *cis*-regulatory fragment of Neurog. Arrows indicate robust ectopic activation of *VAChT -2083/+15>GFP.* The Sox1/2/3 promoter was used to drive Cas9 expression in all CRISPR and control conditions.

**Supplemental Figure 3.**
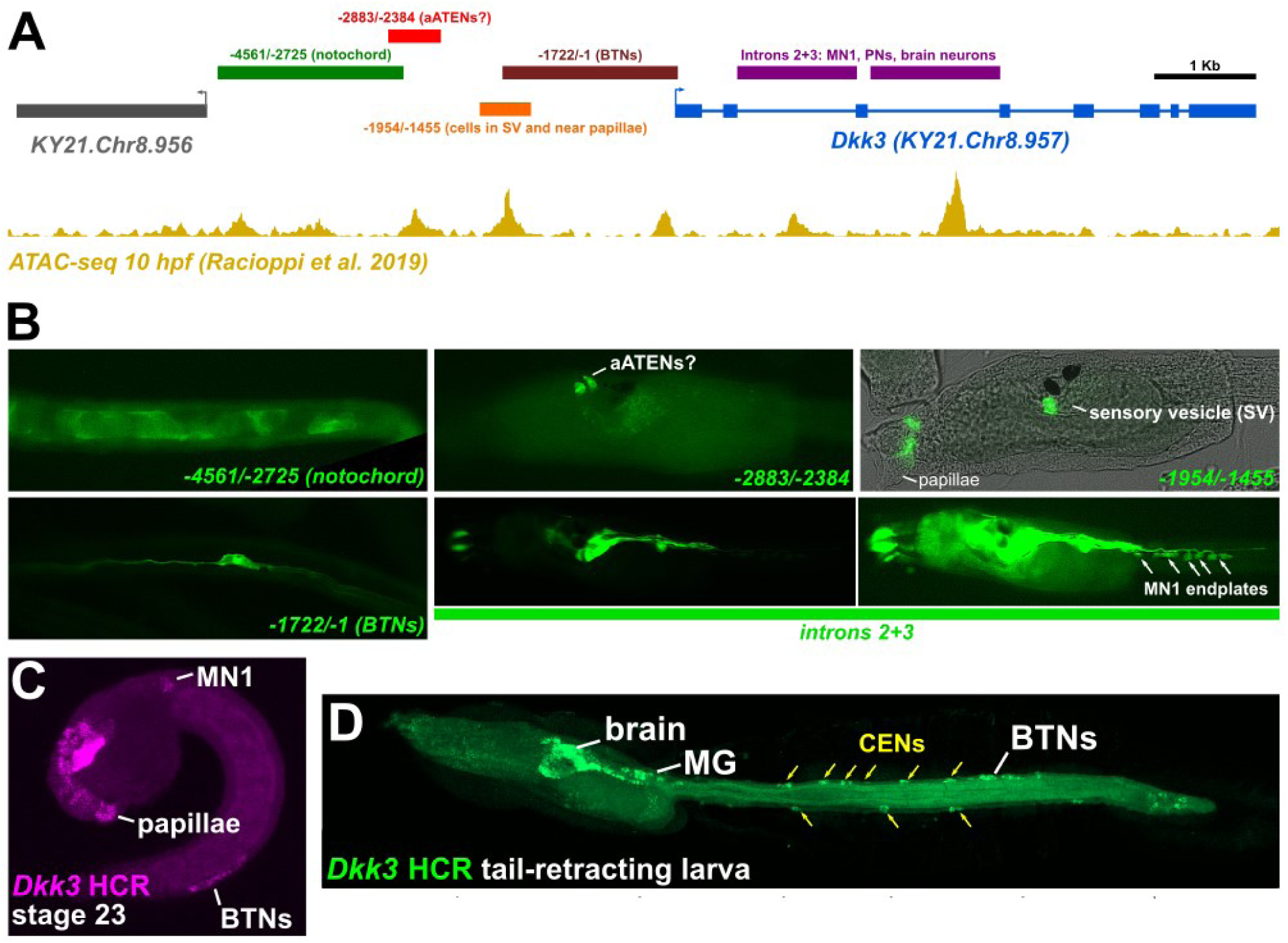
C*i*s*-*regulatory analysis of *Dkk3*. **A)** Diagram showing location and size of candidate *cis-*regulatory fragments tested surrounding the *Dkk3* locus. Ochre peaks represent ATAC-seq peaks observed in “control” (replicate 1) embryos at 10 hours post-fertilization (hpf)(Racioppi et al., 2019). **B)** Representative images of larvae electroporated with *Unc-76::GFP* or *CD4::GFP* reporter plasmids constructed using the fragments depicted at top. aATENs: Anterior Apical Trunk Epidermal Neurons; SV: Sensory Vesicle; BTNs: Bipolar Tail Neurons; MN1: Motor Neuron 1; PNs: Papilla Neurons. Larva electroporated with the introns 2+3 reporter shown at two different levels of fluorescence gain, on the right high gain reveals characteristic endplates of MN1 (arrows). **C)** *In situ* hybridization chain reaction (HCR) staining showing *Dkk3* mRNA expression in MN1, BTNs, and papillae, in a Stage 23 (mid-tailbud) embryo. Expression in papillae and BTNs are recapitulated by certain fragments shown in A and B. **D)** HCR staining showing *Dkk3* expression in a larva undergoing tail retraction just before metamorphosis (Stage ∼31). Yellow arrowheads indicate expression in caudal epidermal neurons (CENs) of both the dorsal and ventral midlines. CEN expression has yet to be confidently recapitulated by any one isolated *cis*-regulatory element.

