## Supplementary material for "Regulation of motor neuron differentiation in the *Ciona* larva": Supplmental Sequences

**Supplemental Sequences File - Popsuj et al.**

**Unpublished single-chain guide RNAs (sgRNAs) used**

Onecut 1.30

**GCTGCATCCGGAGAACTGAA (G+N19)**

Onecut 1.410

**GGAGCTCCAGAAGTCCTCCG (G+N19)**

Lhx3/4 2.45

**GCGACCACCATATATTCGAC (G+N19)**

Lhx3/4 2.75

**GCAAAGTACAGGACAAACCT (G+N19)**

Lhx3/4 3.69

**GCAAGTGGTTAGGAGGGCAC (G+N19)**

Nkx6 1.43

**GGTGTTGAAATGCAGGGTAG (G+N19)**

Nkx6 1.223

**GGATACTTGACGTAAACCGG (G+N19)**

**Previously published sgRNAs**

Neurog.1 (Gandhi et al. 2017)

**GCTTACCATTACGTCTTGTG (G+N19)**

Neurog.p1 (Kim et al. 2020)

**GTGTCTGGCGATAGTATACG (G+N19)**

Neurog.p2 (Kim et al. 2020)

**GACGTAACAAAGCATGACCG (G+N19)**

Ebf.3 (Gandhi et al. 2017)

**GAGACTGTGCCAAGACACCC (G+N19)**

Dkk3.2.100 (Popsuj et al. 2024)

**GTCGATGATTTGGATAAAGA (G+N19)**

Negative Control (Stolfi et al. 2014)

**GCTTTGCTACGATCTACATT (G+N19)**

**Primers for NGS validation of sgRNAs**

| **Gene + exon/sgRNA** | **Forward primer** | **Reverse primer** |
| --- | --- | --- |
| Onecut 1.30 | AAGATTCCCGCTGAGAAATC | CCTTCCAAGCCCATTGATTTC |
| Onecut 1.410 | TTGGATGAAAATTGCAACAAGTC | ACCTCTTTGGGATGATGGAG |
| Lhx3/4 exon 2 | CCCCAACTTACGACATTTATGT | GTCCCCTATAAGAATACCTTGC |
| Lhx3/4 exon 3 | CACCTCAACCAAAATTAACCCT | CCATGACAATTCTACACCAATG |
| Nkx6 exon 1 | GCATCTGCAGGAATGATGGA | CAGTCATCATACCAGGCCAA |

>VAChT -2083/+15 promoter from Hossain et al. 2024, based on Yoshida et al. (see bottom of document for alignment and JASPAR scores)

Exon 1 (non-coding) first coding exon of VAChT start codon ASCI NOTI

ggcgcgccattacgtcgtaaacctttggctaccatcatctgcctcaaaacaaaattaattaaagaaatgcgttagtgtatcctttgactcggaatcaatcaaatcaagcaaaaatcaatatgtgaaattaaccatttagaccttgtgtcattcccattgcggtgacttgtcctagttgtgcgtttttatcagcagattgatttaacagtcaccggaacaggcaagacaatatttcaaaccagccaatgttaatttcagacaaatgaagcaatctgaaaatcagaacaaatcaaaaaacataagttttggtttttaaagcatagaaaacgtaccatattttttaatgtaattgttaaattttgttatttaatatagtagggtagggggagatgggacactttttcatcctgttttctcgtcttggtagcaaacaaaaacattaaaagaattataaaaccgtatcctcgcgactcctacagaccgtttaaaacaggatatttggatattctgttctaaaggtgtcccatcttaccccacagtactatacactatacacattctgtaccgatatattttttgattaaatttgaagttgttaaacttaattacgattaaatttcggcaaattgaaaatgagccatgaattaatcaaaaattattcttgatcgttgttttgtaaaacataactttttttttgatttttttgggaggcgccctgtgttccacttattattatttgtttgttttgctatgtcaatactttaatataaaagaaattaatattggctgacatttcaatttaacgctaggcttatgttttgtttcgtgaataatctgcataaagaaaaaaagcagtatcgactctcctattgttgaatcactcttgcttcccttccattgtgcaggagatagtgacgcaatgaacattgattttaacgcttctagtcagttgggcccttgctcacaattgcctaaaatttgatcaattgtggattgaaaagttaatattctttcctgaattaacatgtctgtaggtaaggtattgagacggctcccatttaatttcttgtccgatctttataaaaaaatatttgttaaaggtacgtttaatttgaacatatagtgcagaagtcttcggaattttcatatagagttattttttagatactatatgcgacatattttccataacccaaattaaattgttttataatttgtttggcttttacagaTTTTAATTTAACTTAGGTGTACGATTCAATAGAAAGATTTTAACCTGTAAAAATACGTGCAAGTTGTTTGGAAATACCTTTTTTGTTGGACAAAGTTGTATCAGgtaagttgatagcatgtatcttaaatctggcgtggtgtttatgtattggcttcatcatgaaatagtttgttttgtccttttacttgtcaattttatttaaattacatgagggtacaattcattataacgtacttcggtgaaaggttaatgttaacaaatgccgtccggtcttctgcgtatctttcgatttcgtattaattatgcatagaaagggtttaagtgcaatgctatttctgatgatgtgtgtgtaacaccaacgacctgatgacgaaaaacttgactgtttattaaataaaatagcgcaacaagacggcggaatttataaatggcatgttgctcgtaaacagcgtgcctgttttaactctggaaaaaaacaagttatattcaactatttgttttcttacgaagcacagtattgcactcttatctaatgtcccaatacaaatatatatatatatatatctagctgttgtcgaggtaaggtgaactgcagttaacaacgaaaaccaacttcgttatacacaaattcaatataaagcgacagctcagtgttgtagagtaggtacccaactttcttaatctgcaaaaggcaaatacatgatttataatatgtaactcagcataacaacctgtttttgccttttgcagATTACATCTGAGGGCGGGTTGTGTGGGTCAGAAATTTATCAGAGGAAAATAAAGTTGATTGAAAGCAAATTTGTTTCTACTTTATTGTTCATCATGGACGTTTGTAGAgcggccgc

>VAChT promoter region from Ciona savignyi from GHOST start codon

(see bottom of document for alignment)

ttaaaagcaaccaacatttcgttcgtaaaactcgtctttctttcattaaaatcaacttggacatttgaattttgaattaaaattccgatcgtttttttaatgcaaaaaagcaactaaatctattttaaaatcgtatttttcaattccaaaattatctaattatagcgcaagttaacatatggctttaaagagcaatagggttacgtactctatcgtctgaggtttttggcgtcttataaataaaaaaatacaatttttattaactgcattcatttaaatttccctgcattattattatgttaaaaatggatataaaaagatcttagttcggtatttgtaaattctgcagaacaattattactggtgacgtattgtaaacgtaaggcttcctttttcgccgctatctcccctaaaattgatttttctcaaaatttctggaccaatgtactttagcctttcgctaattagacattcgggtacagggccgaacactataactcgacgcacgaaaaaggcaactcgacgttagctccattgtctatcgcttcacgtcgacggggatggtagtccaacattgattttatcgcttgagctctatatcgggctcgggttgttaaagatttgatcaatcgcgctaagcaataccggcaacgccaaagtcgttttttttttctaaagcattttgaccgaactgtgttgcattacagcaccagactcagatttggttgtgaaagtgacatatatgatcggttatttgcaggtttttatattttaatagttttatataaagcggtcatagtgttaggcaactttgagtcgttaaagtcgtttttttcacgttttgtgcatccttactagacctgtagtgcattagaccatatcttaatgattaaaataaacacagttatttagtgtaacattgatacgaactattgagatacttataagttcataaaaattaagacgctggttatgtatttttttctgttttaccacattttcttatcttaggttggtagaatttagttatttcaatcataacgcctgattttgattgtgaagaagtaaaatccggacaagcaatttgcgcgttaaatcggtaagtaagcgttgcgtattttcacggggtggatatattcgtgtattttagcatatttaaccaatatggttatcgtaccccctccctgtaaaaaaacaaagcaaaattcatttttgtgtccatttatatcatttgattaccacatatagggtggacagatgtatgttatacaacaacttttgtatttgaatttcgtttccgtattaattatgcatagaaggttttgtgtgcaacgctatttctgatgaattgggtgccacactgtgacccgacacgcaagtaccaccgacgacttaccttgtccaggttatagcttagagggaagtcgagatgcactttggaagcggattttttatagttttaatcaataaagcgcgacgtgcatgaaaccacacgtgggtcgggtgtctgaatacgctttgcgtggtcttgtggattcggtaactttgaatggaaaaggcagtcaattttcatacctttctagtgatgagcccatcatgggttcgaaacttgggccggttaaaaaaagaaatttgttttcagaattataaacggttgtacggccgaatggttaacgcgaacgatatagttaccgatatgtacacaaaaatatcatagtaacaggcgtataggttcgaagcttgggctggtaaaaacgcattgtaatatccaaaacgcataccgaatagatactgtgtgtgaaatagcttagaaaaccggtatttcgttttatttctacaaaaggcaaacacatggcttagtaactacgttaactaaaataataatgttactaaactgtctttgccttttgcagatcgacgaaagaccggtggtctatactcgaaagggaagattttaaagattaaaaaagttgcttctactgagaataccaatatggaaatgttgaaattattcggcaatatgacaatgacatttctgcatcgctgcgttgacggaatg

>VAChT -2083/-742 promoter

Exon 1 (non-coding) ASCI NOTI

ggcgcgccattacgtcgtaaacctttggctaccatcatctgcctcaaaacaaaattaattaaagaaatgcgttagtgtatcctttgactcggaatcaatcaaatcaagcaaaaatcaatatgtgaaattaaccatttagaccttgtgtcattcccattgcggtgacttgtcctagttgtgcgtttttatcagcagattgatttaacagtcaccggaacaggcaagacaatatttcaaaccagccaatgttaatttcagacaaatgaagcaatctgaaaatcagaacaaatcaaaaaacataagttttggtttttaaagcatagaaaacgtaccatattttttaatgtaattgttaaattttgttatttaatatagtagggtagggggagatgggacactttttcatcctgttttctcgtcttggtagcaaacaaaaacattaaaagaattataaaaccgtatcctcgcgactcctacagaccgtttaaaacaggatatttggatattctgttctaaaggtgtcccatcttaccccacagtactatacactatacacattctgtaccgatatattttttgattaaatttgaagttgttaaacttaattacgattaaatttcggcaaattgaaaatgagccatgaattaatcaaaaattattcttgatcgttgttttgtaaaacataactttttttttgatttttttgggaggcgccctgtgttccacttattattatttgtttgttttgctatgtcaatactttaatataaaagaaattaatattggctgacatttcaatttaacgctaggcttatgttttgtttcgtgaataatctgcataaagaaaaaaagcagtatcgactctcctattgttgaatcactcttgcttcccttccattgtgcaggagatagtgacgcaatgaacattgattttaacgcttctagtcagttgggcccttgctcacaattgcctaaaatttgatcaattgtggattgaaaagttaatattctttcctgaattaacatgtctgtaggtaaggtattgagacggctcccatttaatttcttgtccgatctttataaaaaaatatttgttaaaggtacgtttaatttgaacatatagtgcagaagtcttcggaattttcatatagagttattttttagatactatatgcgacatattttccataacccaaattaaattgttttataatttgtttggcttttacagaTTTTAATTTAACTTAGGTGTACGATTCAATAGAAAGATTTTAACCTGTAAAAATACGTGCAAGTTGTTTGGAAATACCTTTTTTGTTGGACAAAGTTGTATCAGgcggccgc

>VAChT -2083/-742 mutated E-boxes (putative Neurog sites)

Exon 1 (non-coding) ASCI NOTI

ggcgcgccattacgtcgtaaacctttggctaccatTGTCCAcctcaaaacaaaattaattaaagaaatgcgttagtgtatcctttgactcggaatcaatcaaatcaagcaaaaatcaatatgtgaaattaaccatttagaccttgtgtcattcccattgcggtgacttgtcctagttgtgcgtttttatcagcagattgatttaacagtcaccggaacaggcaagacaatatttcaaaccagccaatgttaatttcagaTGAACAaagcaatctgaaaatcagaacaaatcaaaaaacataagttttggtttttaaagcatagaaaacgtaccatattttttaatgtaattgttaaattttgttatttaatatagtagggtagggggagatgggacactttttcatcctgttttctcgtcttggtagcaaacaaaaacattaaaagaattataaaaccgtatcctcgcgactcctacagaccgtttaaaacaggatatttggatattctgttctaaaggtgtcccatcttaccccacagtactatacactatacacattctgtaccgatatattttttgattaaatttgaagttgttaaacttaattacgattaaatttcggcaaattgaaaatgagccatgaattaatcaaaaattattcttgatcgttgttttgtaaaacataactttttttttgatttttttgggaggcgccctgtgttccacttattattatttgtttgttttgctatgtcaatactttaatataaaagaaattaatattggctgacatttcaatttaacgctaggcttatgttttgtttcgtgaataatctgcataaagaaaaaaagcagtatcgactctcctattgttgaatcactcttgcttcccttccattgtgcaggagatagtgacgcaatgaacattgattttaacgcttctagtTGGTCAggcccttgctcaTGATCAcctaaaatttgatTGATCAtggattgaaaagttaatattctttcctgaattaacatgtctgtaggtaaggtattgagacggctcccatttaatttcttgtccgatctttataaaaaaatatttgttaaaggtacgtttaatttgaacatatagtgcagaagtcttcggaattttcatatagagttattttttagatactatatgcgacatattttccataacccaaattaaattgttttataatttgtttggcttttacagaTTTTAATTTAACTTAGGTGTACGATTCAATAGAAAGATTTTAACCTGTAAAAATACGTGCAAGTTGTTTGGAAATACCTTTTTTGTTGGACAAAGTTGTATCAGgcggccgc

>VAChT -2083/-742 promoter mutated putative Onecut sites

Exon 1 (non-coding) ASCI NOTI

ggcgcgccattacgtcgtaaacctttggctaccatcatctgcctcaaaacaaaattaattaaagaaatgcgttagtgtatcctttgactcggaatcaatcaaatcaagcaaaaaAAGGAatgtgaaattaaccatttagaccttgtgtcattcccattgcggtgacttgtcctagttgtgcgtttttatcagcagattgatttaacagtcaccggaacaggcaagacaatatttcaaaccagccaatgttaatttcagacaaatgaagcaatctgaaaatcagaacaaatcaaaaaacataagttttggtttttaaagcatagaaaacgtaccatattttttaatgtaattgttaaattttgttatttaatatagtagggtagggggagatgggacactttttcatcctgttttctcgtcttggtagcaaacaaaaacattaaaagaattataaaaccgtatcctcgcgactcctacagaccgtttaaaacaggatatttggatattctgttctaaaggtgtcccatcttaccccacagtactatacactatacacattctgtaccgatatattttttgattaaatttgaagttgttaaacttaattacgattaaatttcggcaaattgaaaatgagccatgaattaatcaaaaattattcttgatcgttgttttgtaaaacataactttttttCCTTtttttttgggaggcgccctgtgttccacttattattatttgtttgttttgctatgtcaatactttaatataaaagaaattaatattggctgacatttcaatttaacgctaggcttatgttttgtttcgtgaataatctgcataaagaaaaaaagcagtatcgactctcctattgttgaatcactcttgcttcccttccattgtgcaggagatagtgacgcaatgaacTCCTTttttaacgcttctagtcagttgggcccttgctcacaattgcctaaaatttgatcaattgtggattgaaaagttaatattctttcctgaattaacatgtctgtaggtaaggtattgagacggctcccatttaatttcttgtccgatctttataaaaaaatatttgttaaaggtacgtttaatttgaacatatagtgcagaagtcttcggaattttcatatagagttattttttagatactatatgcgacatattttccataacccaaattaaattgttttataatttgtttggcttttacagaTTTTAATTTAACTTAGGTGTACGATTCAATAGAAAGATTTTAACCTGTAAAAATACGTGCAAGTTGTTTGGAAATACCTTTTTTGTTGGACAAAGTTGTATCAGgcggccgc

>VAChT -2083/-742 promoter mutated putative Neurog and Onecut sites

Exon 1 (non-coding) ASCI NOTI

ggcgcgccattacgtcgtaaacctttggctaccatTGTCCAcctcaaaacaaaattaattaaagaaatgcgttagtgtatcctttgactcggaatcaatcaaatcaagcaaaaaAAGGAatgtgaaattaaccatttagaccttgtgtcattcccattgcggtgacttgtcctagttgtgcgtttttatcagcagattgatttaacagtcaccggaacaggcaagacaatatttcaaaccagccaatgttaatttcagaTGAACAaagcaatctgaaaatcagaacaaatcaaaaaacataagttttggtttttaaagcatagaaaacgtaccatattttttaatgtaattgttaaattttgttatttaatatagtagggtagggggagatgggacactttttcatcctgttttctcgtcttggtagcaaacaaaaacattaaaagaattataaaaccgtatcctcgcgactcctacagaccgtttaaaacaggatatttggatattctgttctaaaggtgtcccatcttaccccacagtactatacactatacacattctgtaccgatatattttttgattaaatttgaagttgttaaacttaattacgattaaatttcggcaaattgaaaatgagccatgaattaatcaaaaattattcttgatcgttgttttgtaaaacataactttttttCCTTtttttttgggaggcgccctgtgttccacttattattatttgtttgttttgctatgtcaatactttaatataaaagaaattaatattggctgacatttcaatttaacgctaggcttatgttttgtttcgtgaataatctgcataaagaaaaaaagcagtatcgactctcctattgttgaatcactcttgcttcccttccattgtgcaggagatagtgacgcaatgaacTCCTTttttaacgcttctagtTGGTCAggcccttgctcaTGATCAcctaaaatttgatTGATCAtggattgaaaagttaatattctttcctgaattaacatgtctgtaggtaaggtattgagacggctcccatttaatttcttgtccgatctttataaaaaaatatttgttaaaggtacgtttaatttgaacatatagtgcagaagtcttcggaattttcatatagagttattttttagatactatatgcgacatattttccataacccaaattaaattgttttataatttgtttggcttttacagaTTTTAATTTAACTTAGGTGTACGATTCAATAGAAAGATTTTAACCTGTAAAAATACGTGCAAGTTGTTTGGAAATACCTTTTTTGTTGGACAAAGTTGTATCAGgcggccgc

>Sox1/2/3 promoter from Stolfi et al. 2014 ASCI start codon NOTI

GGCGCGCCcgctcgcatgtcaaatagttcgaattttattgtagaactcgggcagttgtgatgtaacacgtgaattggattcagcattcgttctttaaacctatgagtcggcaaggaagcgatctgtcaaattaaatcgaatctttcagcgctcgcatcgagacatagcgatactggaagcgtcggaaatagaatagtggggttgatgacaggtgaaaacagaattcgtggcgcaaattacgaaatatttttgcgctggttttattaaacaacaattttctcaatagtggggtcgaaattgatgtttgcattttgttgtaggggttgtacatgcgggatacgctccgtggtaggcgtctatgttatacagtcggtgtaacttaacgacaaatatagaatacgttgtgttcgttcattgatggtcttgagttgcacgatttcccgtcgcaaatgaaccgttaaacacaacttggcccgaagcaggcatggcagaagtcgtcgtaagtaggctggggtgtcgtggggccgcgcgtcgtcatgctaaaaccgaccgagcagagcgtaggcaaaagtaagcttcgaattgagtaaaacgcgacgaataaaacggtggtacttgtcggtttgtccgcgctgaaacctaataactggctacgcgtgttttctaggatgtcgagtgttcgcggacacttgaacacttagatatctacatgctgtcgttgctttggttggttttacactggatttgtattgctggtgttttgctgtaaagccagagtttgtaaccatatagacatagtatatttgtaatgtagtattcctagttttgagtatggagctcgctgtttagttctgttgtcagttttaagcgtgttaaggcaagatacataaggtaacatagtctttatctatcttgtcgaatagagtaatgaagatgaaattattaaaagtttaaaaacaaggaattaacagcaaaatgacagttggtaaaacacgctatgaataaactgtattgaaaaacatcaggaaatcgtttggtttgtgttcaatttgtttaatttaaaagttttgtattgttagtataatgactgttgtctgtacacaacacgtgttgaccactatagtagtgtagtagcccaataacaattactggtgttctgcacataagtcgcgtgtgctttactgggcattcatgccggtagaattaaccattcatcgtaaaagaacaaaacggtctattctgatccgttcattgagtggcttatgaattggcaaaagtgcttgtaatgtcggcactgaatgagcgctgcctttgttactgacaatgtgcattcatgacgatagttccgaaaagtgtggaagtaaaatacactgtttgtgttcatactacgcattcccaggaggaataccttttagatacgccatgtatgcttgatttttttgactaaattaaacattacaattattggaacgaccataaggttgttagttttacactttgttgatatttttgtatgggtgttatacgttaatgggatgttattgaaatatagagttgtgtaattgtaattgtgcggtttcatgttaacgcttaaaacagttttctatctaactggtgttttgagttttaggtatatatgataaagttttatgtattgggttatgagttttgttatgagttttttctgtaaatataacttttgcgggttttcatatttttcgattcgtataaaatcgaattctgtcttatcatgaaccgaccacgtttttcgctgattcgaagtcgtttagatggttgctttagggaacgctggatcccaacgaaaagaaaaaggcgacgttcgcctccgcgaagatatgccgtagcgagagaaaggtcgtttgtagcgcaataaaggcagcctgtgagatgcctacttcattcattctgccttttgtgacttcatagtggcattgtgagcttctattcaggcattgtctccaagaaagttatgagtctatagattataagacctcttctatggacaagcagaacaattgaattacaattgttaaaaacaatcgtaaattgtagaccgtagtttcaatagaattaacgcgagaatatttctggagtaaaaagactgaaatagaaaaatagggaccattaaaaattgtcagccgatagccagaatattttacaagtcaactgtaatttgtgtttgaagttttttaaattgatttttttaactgagacttctactttacagcgtttggaatcaagtaaagatatttaactcaattcttgcgaacattcgcctaaagtctcacacgtcattaaactggattttgtagcttacaaaaacttctccgtccctactccaccggggtttctgaaagagccatctcagaacgacttcatgGCGGCCGC

>Claudin.j intron 1 + basal promoter of FOG (bpFOG) ASCI XHOI NOTI

GGCGCGCCccaacaatttagtgtggtgtgcaattaaagaggaataaaacattcaagcttacataaactattttgtgtcatattaatttagatcctaaaattatgttttaaaattaccaaaacaatatatcaatcttataaaacgtgtggaagtatattattcaacgtgattgtgtttcagtcagttactctttcatagagcgacgggaaatgaaccgttgaaaatgttccatctgaataatgcatggaccatggctagcttggtaattagcacagagtagtacggcgagtggtactacaatatgcgagtcgtttcgatttgtcaatagacttgctgcataacaactctcattacactcaattaaacaaaatacttttaaaacttcaaaatttactaaacaaatttcactttcgtcgctcaatgagtgtttgttccatctaattggaaaatgtgtcaaaactttacgttatatttgataacaagccctgtactcgcgaattccttttaggtttagacgaaaaatcgatggattttaggtgacagtcggatcaattctcctggttgaataagcaaggcatggacgctaatgaaacttgtcgtatatgtcgccgagttttcacgcgagtgcttatggattaccgataacactgtaatataacagctccaaactcgccaccgatttgttgctggcagttcatattaacatttttttaccgcttatattaatcatgaagcgcaaacgaaattagaagcgttatagtgtttctaaatactttcgttacaattccaaaccattaccacaatgtagtactggatcaataggatatattgtatgtgtgtacaccaacacacatatacagtaaaatagggggaagttgagacgtggggcacgttggaacgaaaacaaattttattcgacagtattcttacagtttaacaaagtgcgcggcgggtttactgttttatacaacgttataggctaaaaaaaaactaaaaaagatggtcgttttgaatactaactgaatcatattaaaactgtcgtttttgtatagtttgctggttagagttatcaatctagtatcatagtttaatacaatgaactgaacgatttcgttcgattttgtctcaagacattatagtttcaaaactgagatcataggttaatatggcttaaattaaagataaaacaagcacaaaaatatcgttttaaaatcatgtgtatatagtagagtgggggaagatgggacacctttaggacccaaaatgtccaaatatcctgatcgtgttttaaacaattaacagcagtctatggtagtcgtgaaaatactttttaaaatttcttttattatatagaataataaggtgtcccatttccccagtatattatataaagtttccgtggcattgaaaccaacacggctcgttaatttcggtaggttaactacaagttgcagataaatattaaaataaaatcttccattttttgttatcccaacatactataggtaacatattaaaactaaggcaatgggacagacaaagttgctttttgaaacgtacaataatattttgtgttttttacgtgctgtagttctaataatgaattggtttattttgtcaaaattaacaaagaatttaagctttcccatcgtaccccgctttttgcgatttgcttaaagtacctggtgtgcgttaaataactccatgtatgttttaatacatttctcagttgttgtcaaatataataatgttacggtagtattcgtggtatcactcaaaaaacgtgatattttttttttttttaaatggtgggcaatatgtgcatctcgtcatgtcccacattgctatatatgtatgcgtgactgcgtgactgctcttaataatttgcacaacccattagtggtcactgcacattgaagcaattactgtaagtgtgtttcccatgcacgcacactcgctcctatctttgatattatgtcaagattaacgtatttagcgaggataacgtttttttcaatgaacacttctctgtatttttatattttctatgcgtagtaaatgtttacaaaacatattcaaaggcatcatttataatgtatacccagtatattcacaattttagataaacataaatttcgagtatggcagttcctctataccgcgtggtttacagacaccacgcggaaatatgtgtctgttCTCGAGtatctgcaggtcgactctagaggatccggcaaagcttcgtgtattgtaccggcccattgtcaatcatgcaaacttgatattatattgacaagagaagaaggcagtttaaattaaaactctaaagtagagagacattaatctcagctgacaaggcaggtggtcacagtaagttcatttaaatagttggccaacaatagcctttccaagaaagtatttttgttccaggtctatacaaaaataacacacataGCGGCCGC

>Fgf8/17/18 -4835/+12 “sec” from Imai et al. 2009

mutation to disrupt endogenous Fgf8/17/18 start codon

gcatagatttaagccacctcgtgaagtgggcgggtttaattaaaacaaacagagctccagcaatcagtgcaatcaagacgtttgtcgtaggttaaaatattgctgctaaagccgcttctgatcagataaattatgataaaatcaaaccgcatttgataacatctgctccgtattgttttgccggtaagtatggacatagacattaaaaaatgtggaataaatatgaaaaacgcggtataatgatggactataggtatatgcctggtttgaaccgaaaatgcaaaggtcacagatctaaagctgttgaacgcaaggtgttcaagcgaaaatttctgtgaaactgctgtttggtatttttccgatgctaattcggtgcgttgagagaaccccatggaatgctgtaatatgctagcgtttgttgttacaataaacatacctaagtttaagcgttataaaaatctagcaatacaccaaacgtacatgtacagaaaagtcgtggtttaccaataataaataagttatgtgtatatatatagcaccggtattctattcttatctcgcaacctcagtttactggactgaaaacaaatcgggcttgtctccctttgttctgagtcttgtccacaacaaagcgtcagatcgcgctggtttaatgatctcgaatttcagatcatcagttttcagtttcccaaatattccaactatccagtaccgctagcacagggtacacaaataggctgtgtatcggctctagcgacttgacctcgtcggtgtttgtgcaaatttataatcccgccagcctaaacagggcaagggccgctgaggccgcattgacatacaagtacgcgatgcttgcaaaatgccagtccgcaatttctcgcaaaaaaacgtgactgcactctcggggtagcgcattgttccggcggctcctgcctgcacggcgttttcatgaatcacgcggctgtctaatctacgatcaacggccttagccaaaaacccggatcggtaaaagttaacactgttgcgaactcaatacctcgaaattcttgaggttactgcggttactcgggccggattatccgtcaaaaataagcgaagtcgggttcagtttacgtttctcctaactattcccaactgaaaatgacaattaaaaaaaggatcatgatattgtgattagcatgtcatccagcaacgagctgtgtatatgtaggagcatgttttccgaaaggattactattaattttgaagtttgatttcagaaaatcgattatgagcaaacaagtaatgtgcggaaacgattataatacctttttacgaagcagtgtaccagttaaaacagtaaatcaaattctgttactaaagcagtcgatgtttaaacggctgggtgtatctcgctttcaagagtcggcgtcggtgtccggaaaccccgcttctcccagaggtagtgttgtgaaaaatgccggcccagctattctcattgtttagcctccgataatgggaaatacaacgcgtgtctgagataccgctttcccattcctcaagtacgttgttcgcctgcctaccgtccattgactgctggttaattacatcaacttgttttatgggaaacgcttatcgcacacagagagcgtgacgcacagaatcagagagcacatgtaatataaagttgtatgtaaacaagggagttttagtccttagtgagcgccatacgattgcgatcggatgtttatcagcctcggctagtgcacagtttaaacgttctgatgttagttgccgagccctctcagcaagtatgtatacagctgtttattcgggcctctgtagttctagtgttaaggaaggcagactcgactgcacgctacaagcgacagcgctttttccatcaatccaatcacatttgttgagccccttacgggcacttgaggtgacggcgctaatttctaagcagtttcgattgtgtgggcagcttggccgcagagcaaagcacagaaaacgcggcgcttcgcattgcagcagcgccgaacactttgccaaggttgcagtatatgaatacagtagcgccgcgaaagacggagagtttttaaagcacaataagcgtgcctgcggtctgtacatacatacagcaaaattttcggcgttgtgtaaagttggtcgcgacaacaatagccctctcggtgtttggtgcttgaagcggcactacaacagcccgctatcataggccgggaattagggcaagtggcggcttagcgagaatttaattttccttgttaagcaaaatcgtgtgaagtgcttctgtcccacgactgttcacttcctgcgacaacgtatagtgagatcgtacttttgtagccaagataccaatatcaataataatattcagctttgcccgctaggttgaggtaaagggtagatattagttgagaatagttgcgtaatatgattaagctcgtgtaccggcttgctttatatatttttgctaaacaaacaagtgtagcgattaaaatcactcaggcctaagccaagggtaaaaccggttaattcggaggttttagaaacagttattttttgcggtttcactaaacttagaataaatacccaagaatattattatggtttataatttttttttaagttttttttttaaatatccattttttggtgattgtgattgatttgtaataagatatttttaacttcgatattaatgtttgcttaaaattacgaaatagagccgtctgctgcgaaggtttgtaaagtgaaacaattagtgggctactgcagttctactttctgtacaaatatacgctttatcacgctgtcaccttgacagacgttattgtaggaaagtttttgtgctgaacaatagccttgtaattagttgcgggggggcctctcggttcaaggccgaatagatacgacctctgatcagcgggtcgaaaacgaacgtaacaatagggaattatttcgtgctgtgacgtcggccgagtggggctgggaaccatttctaaaacattcctggcataatgatacgatgtaagcgcaatatttcacaaacgaaacaagtttacgtgactgtaactgcataagcgttgctttgtcactggaggcgccaaccgcacgataagatatttcagcgacggtcgcaatgaaaacggacaccctattacatgtatgacggccgttaatacctctgtcgctttgacaccttccaattgtcgataacataacccgcaacttgatattctgtgaatattatgacacccggcgccttttgcctgaattgtgtaatcgtgccgcaatattcgaaattgtcctgggtttatttttaaggcagacgtcagagaaattataattctttacatcagatctgataaaccgctcgatttcacggatgtgttgaacaagcgccgtttcaaactgtcgcagttttatttttatcttcgtataaatttacagcggaattcattgaatgctaaactgttgtataaagaagagtgcggaaagacgggccttgttgggacatactatcaaactattctgatcgtgttttaaacaattaacaacggtatatggtattcgcgagaaaacggtcttataatttttgaatattctttgtttactaccgaataggacaagaaaatagaatgaataggtgtcccatctttccccaccctatactatatatgaaccacacgtactccaaacacacttaacagcgtgacatatttcgctattgcgccagtttaaaacacttctgtcttgagttcaataggcatacgtgtcatagttactacacacgttgtaacttttcccgtatcggaagtttttaaaaaaaaataggactctcctatgacgaattatattaattgctcccctggcgtggtttattttgtcttttcccacttattttcctttgcccacgttttatacagttacctttaatttgctacgcgtttgctcgacgggcggcatgagcgttttggggattatgacgtaacaaaacggtttttaatctcttcctctatgttacgacataatagcacctgtggaaaaacacaaacctgttttgttgggaagctgtatggttgtaaagggatgtgtgtttcgttttgttggtctctgctaaatacacgagtgcaatttcagtcaattacgaataccaacgtatgaaaagaatgcgcagattttgctaattggtttgtttaccctcgttcgtgcttggaattctgcaaattgttgttgtgtgggttagaacaatccaatcaccctgaaagttttttttaaatgcagaagaaaaccattaattttgtgcccaaaacaccgtaagctgtagcgagcacaagttcgatatgtgttcatcaaaaatcggcggactttttctcatcgtcgcttttttcccgcggctccagcaacgcgtttgttaaaagcgcgttgcctaccgttatttacattaaacgatgccgatcgctgggcggttcgttatttttactcgtctggtaatggacaaaataacaacccagtcgtgtgcttagattttagcggtatggagcgccacgcgccgaccctcgggaaggacatgctgcacagcgggaaaccgagctttaggatccggcagataattttattcggagtcgatgttagaattgttataagttgttcaggataatagcaaagtgaaagcagaaataaatttaaactttatgctttgtatatttattcagtaagtagttgaataatgtcaattcggatttaatgcattgcgagtataaatagtaaatccgaatatcaaagtgatttctaagggacattattcttctcttggattacatacgaaaatAcgaccctcc

>Dkk3 -1722/-1 promoter ASCI NOTI

ggcgcgcctccaccaatgtaaacgacgaattagcgcaagcttaggtggggccgagcgattagtgtgtgtcagaaacacaccgtcgattgtctcacaacaagtttgtggacggcagtgctggttgaatgcggtaattgattttaaataggtgtggagaagatgcgcttatactgcacgacttatgtaccataactcccaatctcggatatattcgaagttttatattgcgcggttttaaattcttcatacaatgttgtggggtaagatgggataacgttagctcctaaatcccaaatatcttttgatcaggtttaaacaattaacaacgttttcttagagtcgtaaagatgcggttattcaattctgtaaaataatctttgtttactaccaaatggcttggaagagagaatgacaatataatatgtcatgtttcatcttatcgtactatatattaaacgttttctaatcaaattaagttaatatacttgctcaaatttttttaatataatgtaatgaaatatttatgttttttcagctaatttaaaagtaactttccaagtgtaagcccttagcagcaagattatatatgagcaatctcttcgtgatgggtcatgtcgcctctcgcttcgatcatcggcttgtccaagcaaatacaactttatctaagattttaatgttcgattccatacagatcaaaattggatcagggtaaaggctaccatcggtaagtgctgtaagcgctttttgaactaagcttttttacgaaacgaaaggctggagtaaatattctttggtagctgtagcgtaatggtccactgcttaatatttggcggcaaaacggtagcggtgaaaagtaaggatactaaatatttgttctgtatgtttttaattcaattaactagatgtgtttaaatatatttttgacataatacagttatgatgggcgtttttaaacgaaaaatcaccgtattgtggtctggtttcgaaatttgtatgcggatttagacttctttcaaaaatgttccttaattattagttggaaagtttgaatgacgacgacatacgaaatgctgttttaatcgtcagccacaataaagctgtttatatagctttgaccgcaggcagctaaacggcgcccatgttttggggaagcagattaatcccataaaatgtaaatcatgtaactaagttaatttttttaaatcgtgatactgttttcgtatatttttcgcggtttaaagtaaaaagagcgatttgtgtaatcactgagttttcgcaataaatatggttacacggatcggcctaattacgggttatattgatgggatgattttcaagtaaaccgcatttttcacatcgtacctaaaacagcccaaacctgaatttgcagacaattaaaacatagtttcgggaatataagttattgaatctttgagaatttaaaatatttaaaacggtgtgcaaacgtgtggacctcgtgtggcgcgtcttgtataggcgagtctcgtgtttatctcgtggattattttaggaagcgagaaaaaggagacgagagtaaattattaataagaagggagatagattgtccccgcttcggacagctcgtttctacacagcatgcactgtagacttttttggccgcgctgtttatggtttaacagttaacctgcaattgtgttgttttttcagtttagcattagcaagattaaaaatcgtagtaaaaacagaaagcggccgc

>Dkk3 introns 2+3 cis-reg. fragment (to place upstream bpFOG)

ASCI XHOI XBAI intron 2 intron 3

ggcgcgccagatgggtagacgaagttgaggtatactgtcacgtcatatgttttatgattgttttacttatttcccaggaggctagttaaaattgtattagctatatcgttggtgtgtttgtatgtgttataaaaacatattggaaactgctgttaaaattgcacaacactttacgagtgtcatgccagtactatatttcatttatttcatgaaatatttatttaaaaaaccaaggccgtttaaagcaaggttttgtttcgatgtcgtgaatgcatagaactgtgctggggtaagataggacaccgttatggcacctaagtaccatattttctaaccggattttaaacaaaatagaacgattttttagagtcgtggtgctgtggttataattcctgtaaatatagtttttggtaaatgccaaataaggcgagagacgaaaaccatgtcccatcttgccccaccctagtgaattcttttatacgcctcaaaatattcgataattgcatcgggattctgcaattgtcatctgcatggacgtctggtttcatgcatgaaaacacatggttaatacatgtcaacacaatccaaccgctgagattatttccatttgcactcaattttacggtgcgcgcatctgtctcgaatcaataattttcgaaaaaaatcaatacacgaatgccaatatacaataacatcaagagtctaaaggtgttaaaccacatgcctgctttgaagttcgaaattgccttcataaatactaatttcgcgctccacagaataaagttatcgaccaacggactttataggcaaaacaatgttaaacacaaagctgttatgtttcttgcgattaatagataaaattatcgttacatactatgcctaaaaaacctgtgtgttgtttaacacataacccaacgtacaataggttataccagaacgcatcgatgtgtgtaagctatatattttccatatatagggagagagaagatgggacaactttttattctattttctcatttcatttgttagtaaacaaaaattattaaaataattataaaaatgtatcatctttacgactcccacaaactgttgttaattatttaaaacaccataaggatatttgaaaattgtgcgctaaaggtgtcccatatccccccaccctactatatattacctattgtgacattattttcgccccacagCTCGAGcgccgtattacaaaattataattgcattattgttatagatatacggcaagtgggtttcgattttaatgagaacccaccagaaaagcgtgccggaatttagaaagcggtagtcgaacaaattaaaccctgactgtagatatatagtagggtggggtaaactaggacacttttgcattcttttcccgtctcatttttttagtaaacaaagaatagttacagaattatatagcgtagcgcgaaactataaaacaaggttgttaatggttcaaaaatcgatcattaaatatgtgatttagttgcaagtagatcccgtcttacgcccacagtactataattcatttttattgtcgcgtaaaatacgcttaagtaatgcgaaacagcaacgggtttataagcaattaacatttgaaactgaacgtatccctaccattaataaattatttagatagattaggaaagctttttgtcacagcggttccattatttcaattttatgcataatttaaatacgagccaacgtataatttacatggaagaagagttgaaggatttttgtcccatagttttcatacagtcacaccaatgtcatatatgctttcaaccaaaatccaatagttaataatgtcggccgtattaaacagtacgttcaactgattaatattttataaacttgatttgttttgacattaaaatcgcctagaataactacgcctagaaaaatatcgaccgtagcttgacatgtaaacaccatacacccttccataataaagcatggtagggcgaacccgacgccgtggtgatttgtaggaagatatataccgcagagcaattatcgatttccggctctcaacgcaaaataatagccagtctggtcaggcgtacggtctcctgtttaatgagtgtggtgtatacaccagacgctgcggtttctttgtacacacatttactgagacgaggcttgacgtatacactgtttgcttttaacgattaccaatgctttttctgtaaaacctcaaaccggtcctttttatatcatgttttataatggggtccttttaacgtcattgtacagagcgagaaaatgctcacatagcaatattaaacatttttaaggcgagtgacaccatcacgactcccatagaccgttgtacctatttaaaacacgatcatgatatattcggatattacgtgctaagtgtcccatcttccttcaccccaatctattttaaaactgttttattacgtacaataatgagcaagtttctacaatctctgagtaacggtatatgtattactttttgttgtgcaggtcaatctgaatggctgtccgtctaga

>Dkk3 -2883/-2384 cis-reg. fragment (to place upstream bpFOG)

ASCI XHOI

ggcgcgcccttttggtttactacttaatggaacgagaaaacaaaacgaaaagatgtcccatctttccccattctactgtatatacgcttacctatatcataagagcgacagttcaataacagtttataagtcaacaccgcgctacttaatgcaagatacgaaataaatcctccattaaaaattttgcggaagattagtgacgtcagtgcgcgaatgacgtcacaaggtaatctgacacttccgtggcgaaagagtgacaaccaatgataggcggcatttcaatcagctttattggccaatgttattttaaattactgtcaaatattttaaacgctatctctcaataagggtttcctatttctgagaaataaacttccaagcgttttggcatttgtaatgcgctgtctagtgatatagtaggcattaattttcctagattttaagggtatgcagtgtaaatggaaaagtgtacgaccaagtatacgacctaaagctgtgaccacatgtactcgag

>Dkk3 -1954/-1455 cis-reg. fragment (to place upstream bpFOG)

ASCI XHOI

ggcgcgccgatgttaattatacaaaacactatagagaaacatacgatttaaaagctaacgatattcaatgttaccccacagtgcagctttgcagcattaacttccgttgcgtttattttcagggcgtgtaccacattttacatggcagtttctgcaattcaaattcgcttgccaccgtcgtaatccgaatcggattatccggtatatgcctccaccaatgtaaacgacgaattagcgcaagcttaggtggggccgagcgattagtgtgtgtcagaaacacaccgtcgattgtctcacaacaagtttgtggacggcagtgctggttgaatgcggtaattgattttaaataggtgtggagaagatgcgcttatactgcacgacttatgtaccataactcccaatctcggatagattcaaagttttatattgcgcggttttaaattcttcatacaatgttgtggggtaagatgggataacgttagctcctaaatcccaaatatcttttgatctcgag

>Dkk3 -4561/-2725 cis-reg. fragment (to place upstream bpFOG)

ASCI XHOI

gcgcgcccagcatagtctgtttgcacacgaactcttcgcgtgtaattctaagctataataaaaacgcttaatttgacgttggttaaagatttcttgctaaaaggtacgcatgtatactggatatactaatgactctgtctaaactaaagagttgagacaatactaaaaacaaactggctggtatggcagcttcctgccctgtagaacaatgccttatctcgcggtcccacggccaaccgtaaagactcgtagactttcaaagtcaatgcttggtattataggtcccgatcacgtgactcgtttcctcgcatgcattaagtgtacaattccaagctcatatatagtagggtggggtaagatgggacatctctttattgtatttcctcgtaccatttagtagtaaacaaagaacattcaaagaattataaaactgtaacctcacgactccaataaactgtaattaatagttaaaaacacgatcaggatatcatgtgctaaacgggtcccttcttccccccaccctactctatatcaagtgtaaatatcatgtggttgtatttatctcgtaaaacgctaacaccaccacggtcaaagctacagcgctattatactcaaattagcttagcgtatatccatagattagcagcgggacattaacagatttgtacaataataaacaataactggctgcatatatctgttttgtgaacgtttaattatttttatttttttactttactgtattgtctaaatgttagtaatacctgttacactagttgtcattcgccttccttattctatagctgaatttcaaacgtactctatgactgtgatatacactgtatgacaaataacatctcaaactcatagttcgacccgcttcttctacttactttataattccacaaaacgggtcactgcttgaatcctttatcttactagtttttgtgcaaagcaataatagataagaaccaagcgcccgaaatttgtgccggtagcatttaactaacacctactcgctacaaatctcaattactgtttaattcgaaatcttcaagtaaacttaccgctgataagagtttattgctgataaataaacgcctgtagtaagtttacacaaaacacaaagtgcttgcttcgtatagaaataaaagaacgttttttgttaaagtattgaaatagtatacacttggtattgcattcccgtgaaagtttttttttcaatcttttggttgcagagcgcactcgcagattgttttatagttacaaagtgcagcggagtaagttttgtaaacggttggaaaacgaaacagtgtaaagttttatattcgacaactaccgcgggtttgagcgcatattggtctttaaacaaataaatatctgcccactatataagttacgctgttaaatgattgtttgtgtcccgtgtatattttttttaaaaaaatcaggcaaattcaacaattgtgatccgtgggtaaagtaataaaacggattgatgcaaaatacctctactttatttatatatgtatatatgggcaattcaaacttttaaaaatctataataatgtatagatatgtatagtactgtgagagaacatagaacatatttagcacattatatccgattattctgatcgtgttttaaacaattaacaacggtctatggaagtcgtgagacagcgtttttataattctttaaatgcttttggtttactacttaatggaacgagaaaacaaaacgaaaagatgtcccatctttccccattctactgtatatacgcttacctatatcataagagcgacagttcaataacagtttataagtcaacaccgcgctacttagctagctatctcgag

>Fgf8/17/18 -4835/+12 from Imai et al. 2009

(original version with endogenous start codon intact)

gcatagatttaagccacctcgtgaagtgggcgggtttaattaaaacaaacagagctccagcaatcagtgcaatcaagacgtttgtcgtaggttaaaatattgctgctaaagccgcttctgatcagataaattatgataaaatcaaaccgcatttgataacatctgctccgtattgttttgccggtaagtatggacatagacattaaaaaatgtggaataaatatgaaaaacgcggtataatgatggactataggtatatgcctggtttgaaccgaaaatgcaaaggtcacagatctaaagctgttgaacgcaaggtgttcaagcgaaaatttctgtgaaactgctgtttggtatttttccgatgctaattcggtgcgttgagagaaccccatggaatgctgtaatatgctagcgtttgttgttacaataaacatacctaagtttaagcgttataaaaatctagcaatacaccaaacgtacatgtacagaaaagtcgtggtttaccaataataaataagttatgtgtatatatatagcaccggtattctattcttatctcgcaacctcagtttactggactgaaaacaaatcgggcttgtctccctttgttctgagtcttgtccacaacaaagcgtcagatcgcgctggtttaatgatctcgaatttcagatcatcagttttcagtttcccaaatattccaactatccagtaccgctagcacagggtacacaaataggctgtgtatcggctctagcgacttgacctcgtcggtgtttgtgcaaatttataatcccgccagcctaaacagggcaagggccgctgaggccgcattgacatacaagtacgcgatgcttgcaaaatgccagtccgcaatttctcgcaaaaaaacgtgactgcactctcggggtagcgcattgttccggcggctcctgcctgcacggcgttttcatgaatcacgcggctgtctaatctacgatcaacggccttagccaaaaacccggatcggtaaaagttaacactgttgcgaactcaatacctcgaaattcttgaggttactgcggttactcgggccggattatccgtcaaaaataagcgaagtcgggttcagtttacgtttctcctaactattcccaactgaaaatgacaattaaaaaaaggatcatgatattgtgattagcatgtcatccagcaacgagctgtgtatatgtaggagcatgttttccgaaaggattactattaattttgaagtttgatttcagaaaatcgattatgagcaaacaagtaatgtgcggaaacgattataatacctttttacgaagcagtgtaccagttaaaacagtaaatcaaattctgttactaaagcagtcgatgtttaaacggctgggtgtatctcgctttcaagagtcggcgtcggtgtccggaaaccccgcttctcccagaggtagtgttgtgaaaaatgccggcccagctattctcattgtttagcctccgataatgggaaatacaacgcgtgtctgagataccgctttcccattcctcaagtacgttgttcgcctgcctaccgtccattgactgctggttaattacatcaacttgttttatgggaaacgcttatcgcacacagagagcgtgacgcacagaatcagagagcacatgtaatataaagttgtatgtaaacaagggagttttagtccttagtgagcgccatacgattgcgatcggatgtttatcagcctcggctagtgcacagtttaaacgttctgatgttagttgccgagccctctcagcaagtatgtatacagctgtttattcgggcctctgtagttctagtgttaaggaaggcagactcgactgcacgctacaagcgacagcgctttttccatcaatccaatcacatttgttgagccccttacgggcacttgaggtgacggcgctaatttctaagcagtttcgattgtgtgggcagcttggccgcagagcaaagcacagaaaacgcggcgcttcgcattgcagcagcgccgaacactttgccaaggttgcagtatatgaatacagtagcgccgcgaaagacggagagtttttaaagcacaataagcgtgcctgcggtctgtacatacatacagcaaaattttcggcgttgtgtaaagttggtcgcgacaacaatagccctctcggtgtttggtgcttgaagcggcactacaacagcccgctatcataggccgggaattagggcaagtggcggcttagcgagaatttaattttccttgttaagcaaaatcgtgtgaagtgcttctgtcccacgactgttcacttcctgcgacaacgtatagtgagatcgtacttttgtagccaagataccaatatcaataataatattcagctttgcccgctaggttgaggtaaagggtagatattagttgagaatagttgcgtaatatgattaagctcgtgtaccggcttgctttatatatttttgctaaacaaacaagtgtagcgattaaaatcactcaggcctaagccaagggtaaaaccggttaattcggaggttttagaaacagttattttttgcggtttcactaaacttagaataaatacccaagaatattattatggtttataatttttttttaagttttttttttaaatatccattttttggtgattgtgattgatttgtaataagatatttttaacttcgatattaatgtttgcttaaaattacgaaatagagccgtctgctgcgaaggtttgtaaagtgaaacaattagtgggctactgcagttctactttctgtacaaatatacgctttatcacgctgtcaccttgacagacgttattgtaggaaagtttttgtgctgaacaatagccttgtaattagttgcgggggggcctctcggttcaaggccgaatagatacgacctctgatcagcgggtcgaaaacgaacgtaacaatagggaattatttcgtgctgtgacgtcggccgagtggggctgggaaccatttctaaaacattcctggcataatgatacgatgtaagcgcaatatttcacaaacgaaacaagtttacgtgactgtaactgcataagcgttgctttgtcactggaggcgccaaccgcacgataagatatttcagcgacggtcgcaatgaaaacggacaccctattacatgtatgacggccgttaatacctctgtcgctttgacaccttccaattgtcgataacataacccgcaacttgatattctgtgaatattatgacacccggcgccttttgcctgaattgtgtaatcgtgccgcaatattcgaaattgtcctgggtttatttttaaggcagacgtcagagaaattataattctttacatcagatctgataaaccgctcgatttcacggatgtgttgaacaagcgccgtttcaaactgtcgcagttttatttttatcttcgtataaatttacagcggaattcattgaatgctaaactgttgtataaagaagagtgcggaaagacgggccttgttgggacatactatcaaactattctgatcgtgttttaaacaattaacaacggtatatggtattcgcgagaaaacggtcttataatttttgaatattctttgtttactaccgaataggacaagaaaatagaatgaataggtgtcccatctttccccaccctatactatatatgaaccacacgtactccaaacacacttaacagcgtgacatatttcgctattgcgccagtttaaaacacttctgtcttgagttcaataggcatacgtgtcatagttactacacacgttgtaacttttcccgtatcggaagtttttaaaaaaaaataggactctcctatgacgaattatattaattgctcccctggcgtggtttattttgtcttttcccacttattttcctttgcccacgttttatacagttacctttaatttgctacgcgtttgctcgacgggcggcatgagcgttttggggattatgacgtaacaaaacggtttttaatctcttcctctatgttacgacataatagcacctgtggaaaaacacaaacctgttttgttgggaagctgtatggttgtaaagggatgtgtgtttcgttttgttggtctctgctaaatacacgagtgcaatttcagtcaattacgaataccaacgtatgaaaagaatgcgcagattttgctaattggtttgtttaccctcgttcgtgcttggaattctgcaaattgttgttgtgtgggttagaacaatccaatcaccctgaaagttttttttaaatgcagaagaaaaccattaattttgtgcccaaaacaccgtaagctgtagcgagcacaagttcgatatgtgttcatcaaaaatcggcggactttttctcatcgtcgcttttttcccgcggctccagcaacgcgtttgttaaaagcgcgttgcctaccgttatttacattaaacgatgccgatcgctgggcggttcgttatttttactcgtctggtaatggacaaaataacaacccagtcgtgtgcttagattttagcggtatggagcgccacgcgccgaccctcgggaaggacatgctgcacagcgggaaaccgagctttaggatccggcagataattttattcggagtcgatgttagaattgttataagttgttcaggataatagcaaagtgaaagcagaaataaatttaaactttatgctttgtatatttattcagtaagtagttgaataatgtcaattcggatttaatgcattgcgagtataaatagtaaatccgaatatcaaagtgatttctaagggacattattcttctcttggattacatacgaaaatgcgaccctcc

>Mitf intronic fragment + FOG basal promoter, from Abitua et al. 2012

XHOI NOTI ASCI

ggcgcgccctgctaaaacacgctaacagttaaaaacgtatttacatttaattggaagaaacttacatagcaacttaacactgaagttcatgtatttgacttccctctaaatcctgttaagtcgatctgatccaacgctaataacactataggccgcaattccggattagtgttgctttcaacagactaactcggccgcaccgacgttattaattcgatattaaatgcggattcgaaacacgaatacttcctcttagagttgggaatgccttttccccgtacgcgtatttaaatttgccctagagttcgagttagtgaacacaacttcatgacagtatagcagtaatgttcgcattaagttagcatacagctcgcgtagattgcagagatttttaagatggcatttaattacagataacgaaaaataatgaaattactggtagcgtctatatgcagtaaagtgcagtcaaatatatacctagggtgctttaaaaatacagtttattttacgttttttaatctttagatacaaagcatattcgcttagaaaactgttttaacatttttaagttaatttaaaacacgatgcgtgactctgccaaccccgtcgagttctccaaggaaacgtacccacctatccccctaggccccacgcagtacataccccctagctaccgtttcaaacccagtatgggtgataggcaagcccccgaacaccctaaagttggcaagcgggcatggctggatgaatgccaagcactaacgtacagcagaccagatttaaagaagccaaatatatccactaatttaaacgaccccgaaaacgacaactctgcgagacttcggtttacaaaaagtattgtgtatggcgcatattataaatgccaggttgaagaaagagctgcaatggcttataaacagaaggtaaattgggtgcgctgccaagtctcgtaaactagatattcgctcgagtatctgcaggtcgactctagaggatccggcaaagcttcgtgtattgtaccggcccattgtcaatcatgcaaacttgatattatattgacaagagaagaaggcagtttaaattaaaactctaaagtagagagacattaatctcagctgacaaggcaggtggtcacagtaagttcatttaaatagttggccaacaatagcctttccaagaaagtatttttgttccaggtctatacaaaaataacacacaacGCGGCCGC

>AChRA1::GFP from Nishino et al. 2011

ATGatagttttgcgtttactggttatgggtgcgcttgcgtatgtgagcgtggcaaaaacccggtacgatttatcaggagatataatgcaggggtatgacgctaaagtacgaccgagtgacagctacaataactcggtaaaagtcgtgtttaagcttgtgtttaatcaactactagacgtgagcgaagtcaaccaaaaaatcgaaacgaaactgtgggtttatcacaagtggatggaccctcggttaagctgggtgccagaggattacgaaaacctggagtatatatatctacctactaccaacttgtggctgccggagttggttctgtataacaacgccgatggtgactttgctatttctcaatttacgaaagcaaaagtggattatactggaatggtagaatggaaacctccggcaatttttaaaagcttctgcgagattatggtggcagagtttccatttgacacacaaaactgtacgatgaaaattggcccttggtcgcaaggacaagatttactggacatggtaaattcagactgggaagtcaaagaccacatgtgcgagcccccagatgaaacaatgtacgaggaaagcggagaatggttgattcttaaaactggttgctggaagcattacattaaatacgattgctgtcgagggccgtacgtggacatgacttactacttcatattgcaaagacggcctctatatcttgttatcaacattctctttcctacaatgctgttttcgtatctaacctgcgctgtattctatctgccatcggacgctggcgaaaaaataacactcagtatttcgcttctgctttcactgattgtgttcttgctcgttattgttgaagcgattccctcaaccgctaatggagtcccattgctatgccagtacattctatttactatgatattggtttgcctctctattatgataacagtcggtgtattgaatgtacattaccgtgggcctgcaacacatgtcatgtcggatcggatgaaaaagatattcatggtgtggcttccaaaattcatctatagctcgacaatgaaacgattggatccttacaaggaagaaaaaatgttggctggcagacctcccaaaccttacaaagatatttcagatttatctggtcgGGCCCCCATGGTGAGCAAGGGCGAGGAGCTGTTCACCGGGGTGGTGCCCATCCTGGTCGAGCTGGACGGCGACGTAAACGGCCACAAGTTCAGCGTGTCCGGCGAGGGCGAGGGCGATGCCACCTACGGCAAGCTGACCCTGAAGTTCATCTGCACCACCGGCAAGCTGCCCGTGCCCTGGCCCACCCTCGTGACCACCCTGACCTACGGCGTGCAGTGCTTCAGCCGCTACCCCGACCACATGAAGCAGCACGACTTCTTCAAGTCCGCCATGCCCGAAGGCTACGTCCAGGAGCGCACCATCTTCTTCAAGGACGACGGCAACTACAAGACCCGCGCCGAGGTGAAGTTCGAGGGCGACACCCTGGTGAACCGCATCGAGCTGAAGGGCATCGACTTCAAGGAGGACGGCAACATCCTGGGGCACAAGCTGGAGTACAACTACAACAGCCACAACGTCTATATCATGGCCGACAAGCAGAAGAACGGCATCAAGGTGAACTTCAAGATCCGCCACAACATCGAGGACGGCAGCGTGCAGCTCGCCGACCACTACCAGCAGAACACCCCCATCGGCGACGGCCCCGTGCTGCTGCCCGACAACCACTACCTGAGCACCCAGTCCGCCCTGAGCAAAGACCCCAACGAGAAGCGCGATCACATGGTCCTGCTGGAGTTCGTGACCGCCGCCGGGATCACTCTCGGCATGGACGAGCTGTACAAGCGttccccaaaaccagacgaaccacatttaattggtagcgacgttaaaacagctatggacggagtagattatgtgtcagagtgttataaggaccaacgcgaaggacagcagaaagaagacgaatggaaatacgtcgctatggtgttggaccattttctgttgtatatctttatactggcttgcgtggttggcacagtcggtatatttggcaagcggttgcttgaatttatgtcagaacaagagctttttaaaaacctaggcaacgagtgcattctgaaatgccaaagtgattaA

>nls::Cas9::nls from Stolfi et al. 2014

ATGGCTAGCCCCAAAAAGAAGAGGAAAGTGGACAAGAAGTATTCTATCGGACTGGACATCGGGACTAATAGCGTCGGGTGGGCCGTGATCACTGACGAGTACAAGGTGCCCTCTAAGAAGTTCAAGGTGCTCGGGAACACCGACCGGCATTCCATCAAGAAAAATCTGATCGGAGCTCTCCTCTTTGATTCAGGGGAGACCGCTGAAGCAACCCGCCTCAAGCGGACTGCTAGACGGCGGTACACCAGGAGGAAGAACCGGATTTGTTACCTTCAAGAGATATTCTCCAACGAAATGGCAAAGGTCGACGACAGCTTCTTCCATAGGCTGGAAGAATCATTCCTCGTGGAAGAGGATAAGAAGCATGAACGGCATCCCATCTTCGGTAATATCGTCGACGAGGTGGCCTATCACGAGAAATACCCAACCATCTACCATCTTCGCAAAAAGCTGGTGGACTCAACCGACAAGGCAGACCTCCGGCTTATCTACCTGGCCCTGGCCCACATGATCAAGTTCAGAGGCCACTTCCTGATCGAGGGCGACCTCAATCCTGACAATAGCGATGTGGATAAACTGTTCATCCAGCTGGTGCAGACTTACAACCAGCTCTTTGAAGAGAACCCCATCAATGCAAGCGGAGTCGATGCCAAGGCCATTCTGTCAGCCCGGCTGTCAAAGAGCCGCGGACTTGAGAATCTTATCGCTCAGCTGCCGGGTGAAAAGAAAAATGGACTGTTCGGGAACCTGATTGCTCTTTCACTTGGGCTGACTCCCAATTTCAAGTCTAATTTCGACCTGGCAGAGGATGCCAAGCTGCAACTGTCCAAGGACACCTATGATGACGATCTCGACAACCTCCTGGCCCAGATCGGTGACCAATACGCCGACCTTTTCCTTGCTGCTAAGAATCTTTCTGACGCCATCCTGCTGTCTGACATTCTCCGCGTGAACACTGAAATCACCAAGGCCCCTCTTTCAGCTTCAATGATTAAGCGGTATGATGAGCACCACCAGGACCTGACCCTGCTTAAGGCACTCGTCCGGCAGCAGCTTCCGGAGAAGTACAAGGAAATCTTCTTTGACCAGTCAAAGAATGGATACGCCGGCTACATCGACGGAGGTGCCTCCCAAGAGGAATTTTATAAGTTTATCAAACCTATCCTTGAGAAGATGGACGGCACCGAAGAGCTCCTCGTGAAACTGAATCGGGAGGATCTGCTGCGGAAGCAGCGCACTTTCGACAATGGGAGCATTCCCCACCAGATCCATCTTGGGGAGCTTCACGCCATCCTTCGGCGCCAAGAGGACTTCTACCCCTTTCTTAAGGACAACAGGGAGAAGATTGAGAAAATTCTCACTTTCCGCATCCCCTACTACGTGGGACCCCTCGCCAGAGGAAATAGCCGGTTTGCTTGGATGACCAGAAAGTCAGAAGAAACTATCACTCCCTGGAACTTCGAAGAGGTGGTGGACAAGGGAGCCAGCGCTCAGTCATTCATCGAACGGATGACTAACTTCGATAAGAACCTCCCCAATGAGAAGGTCCTGCCGAAACATTCCCTGCTCTACGAGTACTTTACCGTGTACAACGAGCTGACCAAGGTGAAATATGTCACCGAAGGGATGAGGAAGCCCGCATTCCTGTCAGGCGAACAAAAGAAGGCAATTGTGGACCTTCTGTTCAAGACCAATAGAAAGGTGACCGTGAAGCAGCTGAAGGAGGACTATTTCAAGAAAATTGAATGCTTCGACTCTGTGGAGATTAGCGGGGTCGAAGATCGGTTCAACGCAAGCCTGGGTACCTACCATGATCTGCTTAAGATCATCAAGGACAAGGATTTTCTGGACAATGAGGAGAACGAGGACATCCTTGAGGACATTGTCCTGACTCTCACTCTGTTCGAGGACCGGGAAATGATCGAGGAGAGGCTTAAGACCTACGCCCATCTGTTCGACGATAAAGTGATGAAGCAACTTAAACGGAGAAGATATACCGGATGGGGACGCCTTAGCCGCAAACTCATCAACGGAATCCGGGACAAACAGAGCGGAAAGACCATTCTTGATTTCCTTAAGAGCGACGGATTCGCTAATCGCAACTTCATGCAACTTATCCATGATGATTCCCTGACCTTTAAGGAGGACATCCAGAAGGCCCAAGTGTCTGGACAAGGTGACTCACTGCACGAGCATATCGCAAATCTGGCTGGTTCACCCGCTATTAAGAAGGGTATTCTCCAGACCGTGAAAGTCGTGGACGAGCTGGTCAAGGTGATGGGTCGCCATAAACCAGAGAACATTGTCATCGAGATGGCCAGGGAAAACCAGACTACCCAGAAGGGACAGAAGAACAGCAGGGAGCGGATGAAAAGAATTGAGGAAGGGATTAAGGAGCTCGGGTCACAGATCCTTAAAGAGCACCCGGTGGAAAACACCCAGCTTCAGAATGAGAAGCTCTATCTGTACTACCTTCAAAATGGACGCGATATGTATGTGGACCAAGAGCTTGATATCAACAGGCTCTCAGACTACGACGTGGACCACATCGTCCCTCAGAGCTTCCTCAAAGACGACTCAATTGACAATAAGGTGCTGACTCGCTCAGACAAGAACCGGGGAAAGTCAGATAACGTGCCCTCAGAGGAAGTCGTGAAAAAGATGAAGAACTATTGGCGCCAGCTTCTGAACGCAAAGCTGATCACTCAGCGGAAGTTCGACAATCTCACTAAGGCTGAGAGGGGCGGACTGAGCGAACTGGACAAAGCAGGATTCATTAAACGGCAACTTGTGGAGACTCGGCAGATTACTAAACATGTCGCCCAAATCCTTGACTCACGCATGAATACCAAGTACGACGAAAACGACAAACTTATCCGCGAGGTGAAGGTGATTACCCTGAAGTCCAAGCTGGTCAGCGATTTCAGAAAGGACTTTCAATTCTACAAAGTGCGGGAGATCAATAACTATCATCATGCTCATGACGCATATCTGAATGCCGTGGTGGGAACCGCCCTGATCAAGAAGTACCCAAAGCTGGAAAGCGAGTTCGTGTACGGAGACTACAAGGTCTACGACGTGCGCAAGATGATTGCCAAATCTGAGCAGGAGATCGGAAAGGCCACCGCAAAGTACTTCTTCTACAGCAACATCATGAATTTCTTCAAGACCGAAATCACCCTTGCAAACGGTGAGATCCGGAAGAGGCCGCTCATCGAGACTAATGGGGAGACTGGCGAAATCGTGTGGGACAAGGGCAGAGATTTCGCTACCGTGCGCAAAGTGCTTTCTATGCCTCAAGTGAACATCGTGAAGAAAACCGAGGTGCAAACCGGAGGCTTTTCTAAGGAATCAATCCTCCCCAAGCGCAACTCCGACAAGCTCATTGCAAGGAAGAAGGATTGGGACCCTAAGAAGTACGGCGGATTCGATTCACCAACTGTGGCTTATTCTGTCCTGGTCGTGGCTAAGGTGGAAAAAGGAAAGTCTAAGAAGCTCAAGAGCGTGAAGGAACTGCTGGGTATCACCATTATGGAGCGCAGCTCCTTCGAGAAGAACCCAATTGACTTTCTCGAAGCCAAAGGTTACAAGGAAGTCAAGAAGGACCTTATCATCAAGCTCCCAAAGTATAGCCTGTTCGAACTGGAGAATGGGCGGAAGCGGATGCTCGCCTCCGCTGGCGAACTTCAGAAGGGTAATGAGCTGGCTCTCCCCTCCAAGTACGTGAATTTCCTCTACCTTGCAAGCCATTACGAGAAGCTGAAGGGGAGCCCCGAGGACAACGAGCAAAAGCAACTGTTTGTGGAGCAGCATAAGCATTATCTGGACGAGATCATTGAGCAGATTTCCGAGTTTTCTAAACGCGTCATTCTCGCTGATGCCAACCTCGATAAAGTCCTTAGCGCATACAATAAGCACAGAGACAAACCAATTCGGGAGCAGGCTGAGAATATCATCCACCTGTTCACCCTCACCAATCTTGGTGCCCCTGCCGCATTCAAGTACTTCGACACCACCATCGACCGGAAACGCTATACCTCCACCAAAGAAGTGCTGGACGCCACCCTCATCCACCAGAGCATCACCGGACTTTACGAAACTCGGATTGACCTCTCACAGCTCGGAGGGGATGAGGGAGCTCCCAAGAAAAAGCGCAAGGTAGGTTAA

>nls::Cas9::nls::CionaGeminin-Nterminus (from Song et al. 2022)

nls::Cas9::nls (described in Stolfi et al. 2014)

Ciona robusta Geminin N-terminus

ATGGCTAGCCCCAAAAAGAAGAGGAAAGTGGACAAGAAGTATTCTATCGGACTGGACATCGGGACTAATAGCGTCGGGTGGGCCGTGATCACTGACGAGTACAAGGTGCCCTCTAAGAAGTTCAAGGTGCTCGGGAACACCGACCGGCATTCCATCAAGAAAAATCTGATCGGAGCTCTCCTCTTTGATTCAGGGGAGACCGCTGAAGCAACCCGCCTCAAGCGGACTGCTAGACGGCGGTACACCAGGAGGAAGAACCGGATTTGTTACCTTCAAGAGATATTCTCCAACGAAATGGCAAAGGTCGACGACAGCTTCTTCCATAGGCTGGAAGAATCATTCCTCGTGGAAGAGGATAAGAAGCATGAACGGCATCCCATCTTCGGTAATATCGTCGACGAGGTGGCCTATCACGAGAAATACCCAACCATCTACCATCTTCGCAAAAAGCTGGTGGACTCAACCGACAAGGCAGACCTCCGGCTTATCTACCTGGCCCTGGCCCACATGATCAAGTTCAGAGGCCACTTCCTGATCGAGGGCGACCTCAATCCTGACAATAGCGATGTGGATAAACTGTTCATCCAGCTGGTGCAGACTTACAACCAGCTCTTTGAAGAGAACCCCATCAATGCAAGCGGAGTCGATGCCAAGGCCATTCTGTCAGCCCGGCTGTCAAAGAGCCGCAGACTTGAGAATCTTATCGCTCAGCTGCCGGGTGAAAAGAAAAATGGACTGTTCGGGAACCTGATTGCTCTTTCACTTGGGCTGACTCCCAATTTCAAGTCTAATTTCGACCTGGCAGAGGATGCCAAGCTGCAACTGTCCAAGGACACCTATGATGACGATCTCGACAACCTCCTGGCCCAGATCGGTGACCAATACGCCGACCTTTTCCTTGCTGCTAAGAATCTTTCTGACGCCATCCTGCTGTCTGACATTCTCCGCGTGAACACTGAAATCACCAAGGCCCCTCTTTCAGCTTCAATGATTAAGCGGTATGATGAGCACCACCAGGACCTGACCCTGCTTAAGGCACTCGTCCGGCAGCAGCTTCCGGAGAAGTACAAGGAAATCTTCTTTGACCAGTCAAAGAATGGATACGCCGGCTACATCGACGGAGGTGCCTCCCAAGAGGAATTTTATAAGTTTATCAAACCTATCCTTGAGAAGATGGACGGCACCGAAGAGCTCCTCGTGAAACTGAATCGGGAGGATCTGCTGCGGAAGCAGCGCACTTTCGACAATGGGAGCATTCCCCACCAGATCCATCTTGGGGAGCTTCACGCCATCCTTCGGCGCCAAGAGGACTTCTACCCCTTTCTTAAGGACAACAGGGAGAAGATTGAGAAAATTCTCACTTTCCGCATCCCCTACTACGTGGGACCCCTCGCCAGAGGAAATAGCCGGTTTGCTTGGATGACCAGAAAGTCAGAAGAAACTATCACTCCCTGGAACTTCGAAGAGGTGGTGGACAAGGGAGCCAGCGCTCAGTCATTCATCGAACGGATGACTAACTTCGATAAGAACCTCCCCAATGAGAAGGTCCTGCCGAAACATTCCCTGCTCTACGAGTACTTTACCGTGTACAACGAGCTGACCAAGGTGAAATATGTCACCGAAGGGATGAGGAAGCCCGCATTCCTGTCAGGCGAACAAAAGAAGGCAATTGTGGACCTTCTGTTCAAGACCAATAGAAAGGTGACCGTGAAGCAGCTGAAGGAGGACTATTTCAAGAAAATTGAATGCTTCGACTCTGTGGAGATTAGCGGGGTCGAAGATCGGTTCAACGCAAGCCTGGGTACCTACCATGATCTGCTTAAGATCATCAAGGACAAGGATTTTCTGGACAATGAGGAGAACGAGGACATCCTTGAGGACATTGTCCTGACTCTCACTCTGTTCGAGGACCGGGAAATGATCGAGGAGAGGCTTAAGACCTACGCCCATCTGTTCGACGATAAAGTGATGAAGCAACTTAAACGGAGAAGATATACCGGATGGGGACGCCTTAGCCGCAAACTCATCAACGGAATCCGGGACAAACAGAGCGGAAAGACCATTCTTGATTTCCTTAAGAGCGACGGATTCGCTAATCGCAACTTCATGCAACTTATCCATGATGATTCCCTGACCTTTAAGGAGGACATCCAGAAGGCCCAAGTGTCTGGACAAGGTGACTCACTGCACGAGCATATCGCAAATCTGGCTGGTTCACCCGCTATTAAGAAGGGTATTCTCCAGACCGTGAAAGTCGTGGACGAGCTGGTCAAGGTGATGGGTCGCCATAAACCAGAGAACATTGTCATCGAGATGGCCAGGGAAAACCAGACTACCCAGAAGGGACAGAAGAACAGCAGGGAGCGGATGAAAAGAATTGAGGAAGGGATTAAGGAGCTCGGGTCACAGATCCTTAAAGAGCACCCGGTGGAAAACACCCAGCTTCAGAATGAGAAGCTCTATCTGTACTACCTTCAAAATGGACGCGATATGTATGTGGACCAAGAGCTTGATATCAACAGGCTCTCAGACTACGACGTGGACCACATCGTCCCTCAGAGCTTCCTCAAAGACGACTCAATTGACAATAAGGTGCTGACTCGCTCAGACAAGAACCGGGGAAAGTCAGATAACGTGCCCTCAGAGGAAGTCGTGAAAAAGATGAAGAACTATTGGCGCCAGCTTCTGAACGCAAAGCTGATCACTCAGCGGAAGTTCGACAATCTCACTAAGGCTGAGAGGGGCGGACTGAGCGAACTGGACAAAGCAGGATTCATTAAACGGCAACTTGTGGAGACTCGGCAGATTACTAAACATGTCGCCCAAATCCTTGACTCACGCATGAATACCAAGTACGACGAAAACGACAAACTTATCCGCGAGGTGAAGGTGATTACCCTGAAGTCCAAGCTGGTCAGCGATTTCAGAAAGGACTTTCAATTCTACAAAGTGCGGGAGATCAATAACTATCATCATGCTCATGACGCATATCTGAATGCCGTGGTGGGAACCGCCCTGATCAAGAAGTACCCAAAGCTGGAAAGCGAGTTCGTGTACGGAGACTACAAGGTCTACGACGTGCGCAAGATGATTGCCAAATCTGAGCAGGAGATCGGAAAGGCCACCGCAAAGTACTTCTTCTACAGCAACATCATGAATTTCTTCAAGACCGAAATCACCCTTGCAAACGGTGAGATCCGGAAGAGGCCGCTCATCGAGACTAATGGGGAGACTGGCGAAATCGTGTGGGACAAGGGCAGAGATTTCGCTACCGTGCGCAAAGTGCTTTCTATGCCTCAAGTGAACATCGTGAAGAAAACCGAGGTGCAAACCGGAGGCTTTTCTAAGGAATCAATCCTCCCCAAGCGCAACTCCGACAAGCTCATTGCAAGGAAGAAGGATTGGGACCCTAAGAAGTACGGCGGATTCGATTCACCAACTGTGGCTTATTCTGTCCTGGTCGTGGCTAAGGTGGAAAAAGGAAAGTCTAAGAAGCTCAAGAGCGTGAAGGAACTGCTGGGTATCACCATTATGGAGCGCAGCTCCTTCGAGAAGAACCCAATTGACTTTCTCGAAGCCAAAGGTTACAAGGAAGTCAAGAAGGACCTTATCATCAAGCTCCCAAAGTATAGCCTGTTCGAACTGGAGAATGGGCGGAAGCGGATGCTCGCCTCCGCTGGCGAACTTCAGAAGGGTAATGAGCTGGCTCTCCCCTCCAAGTACGTGAATTTCCTCTACCTTGCAAGCCATTACGAGAAGCTGAAGGGGAGCCCCGAGGACAACGAGCAAAAGCAACTGTTTGTGGAGCAGCATAAGCATTATCTGGACGAGATCATTGAGCAGATTTCCGAGTTTTCTAAACGCGTCATTCTCGCTGATGCCAACCTCGATAAAGTCCTTAGCGCATACAATAAGCACAGAGACAAACCAATTCGGGAGCAGGCTGAGAATATCATCCACCTGTTCACCCTCACCAATCTTGGTGCCCCTGCCGCATTCAAGTACTTCGACACCACCATCGACCGGAAACGCTATACCTCCACCAAAGAAGTGCTGGACGCCACCCTCATCCACCAGAGCATCACCGGACTTTACGAAACTCGGATTGACCTCTCACAGCTCGGAGGGGATGAGGGAGCTCCCAAGAAAAAGCGCAAGGTAATGGCCACGAAAAATATTCTTCAAAATATAAATGCACAATGGAAGGAGAATGACAACAGATCACCAAGTAGAAAGCGACGGTTAGATGACGTCACTGAAGAATCACAATTACCTTCCACGACCAAACGACGTCATCTTCAAACAAATACAAACGTTGTAAATTCCACAGGATTGAAACAAGGCCTGACAAATGTGAAAAATTCAATAAATCCAAAGAACAAATCAATAAAAAATTTCTTTTCTGATATTCCACGTGTGTCATGTACTAAATCTGAAAAGATTCAAATTTTTAAAGAAGCTAAGAAAACTCCAAAAAAGAATGCAACCACTCAGACAAGGAGTGAAGCTGAAGAATTGGTCTGCAGTGATCAACCCAGTGAAAAATATTGGGAACTCTTAGCCGAGGAGCGAAGGAAAGGGTTGTAA

>Stabilized beta catenin based on Wada et al. 2008, Abitua et al. 2012

atgaacttccccgatacaaaccaagtactcggggaatggcaacgagagtttacaatggaagaagcaaaccaaatccatgatcagtttgtgcgcacacgtgccgatcgtgtccgcgacgtgctgttccctgaaaccatgatggaggaaagtgggcccgtcccaagcacacaatacgagtcgaacacggcaacatccgtgcagagattggcagaaccatcgcaacaacttaagaaggcagtggtaaacttgatcaattatcaggacgatgccgacctcgccacgaaagcgatccccgagttgactggtcttctcaatgatgacgaccaagtggttgttcaacaagcagcacaaatggtgcacatgttgtccaagaaagttgcgagtagacaagcgataatgaatagcccggccatggtgtcggctctcgttagagccatgcagaacgccaccgaccccgagacccagcgttattgtaccggcgccctacacaacttgtcgcatcacaaacaaggattgttgtcaatattcaaatccgggggaattcctgctcttgtcaaaatgctcagttccccaattgagtcggttgtgttctacgcgatcaccaccctgcataacctacttttgcaccaagagggagctaaagaggccgtgagattggcaggggggttgcagaaaatggtgtatctattgtcgcgtgacaacgtaaagtttctggcaattgacaccgattgtttacgaatcctagcttatggaaatcaagagagcaagttgatcattcttgcgagcaacggaccccaggaattggtccgcatcatgaggacttacgattatgagaagttattatggacgactagtcgcgtgatcaaggtgttgtcggtttgttcgagtaacaaaccagccatcgtagaagcaggtggaatgcaagcacttggcatgcaccttgggtcaccctcacaaaggctgttacaaaactgtttgtggagtcttcgaaacttatctgatgctggaaccaaacaagatcatgtggagaacttgctacagatgcttgtccagcttctatcatcgaatgacatcaacgtggttacatgcgctgcggggattctaagcaaccttacttgcaacaacatgagtaataagaccagagtgtgtcaggttggaggaatcgaagctttggtgcgcactgtgttgcaagctggtgatcgtgaggatatcacggaacccacagtgtgtgcgttgcgtcatcttacttcgcgtcacccagacgctgagatggcacaaaatgctgttcggttgcattacggcctcccagtgttggtcaaactgcttcacccaccttcaagatggccgctcattaaggctgttgttggtttgattagaaacttggctttgtgcgctgccaaccacgcggcacttcgtgagcatggggcgataccacggcttgtgcaactactaatgcgtgcccaccaagacacacagcgtaggacaagcatggcgtcgagtcatagccagatgtctgctgcttatgtggacggagtacgaatggaggaaatagtggaaggaacaactggaacactccatatattggctcgtgaacctcacagtagatcggtcatccgaggacttaacacaatccctctgtttgtgcaacttctttattcacaggttgagaacatccagcgggtggcagcaggcgtgttgtgcgaattagcgcaagacagggaaagtgccgacctcatcgagaacgagggggcgagcgctccattgaccgagttgttgcattccaaaaatgaaggagttgccacgtacgcggctgctgcattattccgcatgtccgaggataaatcgcaagattacaagaagagattgtctgttgagttgacgagctccttgttcaaggatgatggagctttatataatggtgcggacctaatggaccccggcatgcaaggttaccaacaccagatgacccccccgatgtctcatcaaagctccgtcagttcagttcataactcgcaccattctgggttcccacaagcagaaggccaacaacccatggactttcaacaacacctcgtccccccggcagattacccgctaccagacctttcaaacgacttggaccttgaaccaatgattggtgttcaacaaccatggctcgatactgacctctaa

>TCF^Nter^ based on Kaplan et al 2019 (nuclear localization signal)

atgcctcagttaaactcggatgaagctgccaatgatgagccaaaaacctataacgatgaaagggtaggcgaagaggacgaacgagggtggcatgaaaatgatgaacttgacaacataaagggtgatttagtcgaggaagaagacagagacccacacaggcgacaatcttacgatgttccggcaaaagtaaactctcgacacagagattccagccacactagtgtgagcaagggcgaggaggataacatggccatcatcaaggaaaaaaagaaaagaaaagttTAATGA

>Onecut in situ probe template

atggcaaccgctgcatccggagaactgaacggtttccaccatctgcatcatcaccaccatccatcggagcaatattacaggcacgagcactatcaccatcacttccaccaccccaactttgacggttaccccaactacaacgatagagatacccctatcgctggagacatgcagaaaaacaacctgcataatttcgcatcgaaatcaatgagcttggaaggtgaaaaattggatgaaaattgcaacaagtctcctaactatcttccgcctatcggagacgctcttcttcgtcgtgacaatcgttccgatgcctcgaagaacaatgcaaaggaagaagatgagagcgggtgctcaaaattcgtgatgcaagagaccgacaactctctcacagagctccagaagtcctccgcggtcagcgagcacgagaaaaaagaagaagtgcagttgaaaactaacgatgcgccggaagatttttcagtgaagactgagcaatcagagttgtatcaattccacgcgaggaacttctcaatctttactccatcatcccaaagaggtactccggatgaaggaatgaatcttatccccgttgagaccacggaccacacatcaatagattcttacttcagatccgatgcaaccaatgccaaccctaacagcaacccaatagacagcgtcccttccagcgtcgacgggccaagttacgctactctcacacccctacaacctctgccatcgatatcatcagtttccgacaaatacatgcccaccaatgaaaccagctacgcaactcttaccaaccaagagcttaccgattgcagcagctactcaaaaatgggcggaatggggcacagtttacctccgctttcaaacagaatgatcttgaacggtcttgcagctcaaactcgtggaggaatgcaatcacaagctgctatagatgcggttaaccaagcggcagctgcagcagttgggctttcacattacaacaaaccagtgttgtcatctaatataataccaccaccacctccggtttcaaacccctacgatccccatgttttcgggaggattgatcaatgcaatgatatgggtgctggctttcctggcgggcatatgtttccgcatagaagtactgggtttgtatcacaatatgggctccaagacctatcttcatcactccaagtcagcgcgccttctgagcgaagaagaccgactcacgaagatattcccgccgacaatggaaagcgccattccggcagcgatcgcctcggagggtcgggcctgcagccgcattcatctaactccgctagttcttcccgcactcaacagatcgaagaagtcaacacgaaagaggttgcttcgaaaatcacccaagaacttaaacgttatagcatcccgcaagcaatttttgcacaaagggtgctttgtagaagccagggcacactttcggatcttctacgcaatcctaaaccgtggtcaaaactgaagagtggacgcgagaccttccgaaggatgtggaaatggcttcaggaaccagagtttcaacgtatgtcatctttaaggcttgccgcttgcaaacgaaaagaggacgaaaaatcttacgagaactcggtcaattctccgaagaaacctcgcttagttttcacagatctgcagcgtagaactttacacgctattttcaaagaaagcaaacgcccttctaaagaaatgcagatacaaatctctcaacagttgggtttagaagtcacaactgtgagcaatttctttatgaacgctcgaaggagaagtttagataaatggcaggacgacgaatcagggtataacagcaaggaaaactcacgttctaacaacccatcaagcgatcaccatctaagtgcgtcgcctaaccaccaacaacagcagcagcaacaacaacaacaacagcaacaacagcagcaacaagcgtatcaacagcatacacaagacagtcgtttaagttatgcgccaggagagtctcttctttccccgctctgtggctcccctagcggccatttgcacttcccaccccctcaccatttgcaccaccataacttgcaccagcagcagcaaaacacaatgttatcagcgtcacacctaacctcgtccggtctagtacacccgtaccagtcacagcaccagcttctaggctctgacgtcacggggttggtgaatccccgttag

>Dkk3 in situ probe template

atgaaacgactcgtcgtcttttctttcattctgctggtgaatttttgttcggggatgtatcggtataggttgggcggagagctggaccgtcaactgtctccctacaccattgaaaacgatgctaatgaccgagaggaggaggaaagggaagatatgttgtataaatacatgatggagcagcttattcgtgatataggaaaggaaaatacctttgaaaacgaactccaagacaacgatagtccccccagtgctttagaagagctgactcgtcttgagaacacaatcgatgatttggataaagacggaagatgggtagacgaagttgagacgctggaaaacgaatttcctcctaactaccacaacgtcacagagtttcgaaccaaaattgggaatcagtcagttcatgtgaaagaatctatcgacaaaaacaccctggggaatgaaggtcaatctgaatggctgtccgaagaaatcacctcaaaaagcgatgtcaaacgagaccgaggcccatggaaatgttcgatcgcattcccatgttcagaagacgaatattgctttgaaactcaaacatcaagcgagtgcaaagaatgtaaattagaaggagaccaatgtgtggataatgcggagtgttgtcaggaacctcatgccaactacgatcaccaagctatgtgtgtattcggtcggtgtaaactatctacaagaccaggagcaattggtaccatttgtgaggttcctgacgattgtgatgctggtatgtgttgtgcggacacacctggctacgcgagatcattatgtaaaccaaacgccagggttggagagcgttgctcggttgaaccagattacaatccatctgaatacattgtctttgaagatgggcaatatttttgtccatgcgaatcggacctgacttgtgttagcagaggtacaaacaaagaggacaacctacaggattataacactccacggttctgcgaaatgcctcggctaaacttagatcgtgcgagtacaaaggacaaggtgcaaattgaaagagaccttttagcaaagctgtttccggaactggttgtaccgcactggcagcagcgtatggagacgatagaagaccaacagttcacttcaaaaatgaatccaagtcttcacaaaacgaccaaaaaagcaatggaggaaaacgaattgatcatgtaa

>Lhx3/4 in situ probe template (Stolfi et al. 2011)

agacaatgcagaccggaagtgagtttcatcaaaattcagccggaagtatacgtcaccaaccggatattgcgtaccaccaagacagaaagcatttatcgacgcaacaatgggatgacaatggtaaaaaatccgacgtgaatcaagatgacgattttgaccaatacgtggacgatgacgtaggttatgacgtaggcgatgattttgatgatgacgacgacgatgacggcatagtggtcgattgtgatgacgacagagactcgctgatggatttagatttttcaacatccttgctttccggtcacagtggggatttccaggatcgaaaacacgactttcctacgtcatcgctgcaagaactattctctatgacgcaatcggtgacgtcatcaagttgtgacgtcaccaacctcgtgaccagctccatccaatcacatcccgacctcaacgactcggggatcgttaccacagtaggggaatccccgcagaacccctttttaccccaaacctcttcgaaacaaacggacagcgctgggtcgctgttcgcgcttctatcgtccgatcaacgaatacaagccaagataccaaaatgcaccggatgcgatcaccatatattcgaccggtatattctcaaagtacaggacaaaccttggcattcacaatgtttaaaatgcaacgactgtgggcgacaactgaccgataaatgtttttctcgcggtagttacgtttactgcaaagaagatttcttcaagaggttcggcaccaaatgctccggttgcgaactcgccattccccccacacaagtggttaggagggcacaggacaacgtgtaccatctagaatgcttcagatgtttcatgtgtagtgagcaattgggcaccggggaccaattctatttgctggatgatagtcggcttgtgtgtaagaaagattacgaacatgcaaagtcaagagatcttgacatggacaacggtataaagcgaccccgcaccactattacagctaagcaactggaaacactgaagatcgcttacaatcaaagcccgaaacctgctcgccacgtgagggagcagctgagctcggacaccgggttagacatgagagtagttcaggtatggtttcaaaacagaagggcgaaagaaaaacgtttaaagaaagacacaggccgacaacgttggggagagctattccgatcaggggcgccgtctagtggccctcattgtcgaccaaacccagactccccgcccagcggtggtaagcgtcgtgttggtggccatagcaaccgtaagcggccctctagctctcctggtggcacacgtatagctatcccgataccatcgtcgattcaaagccctggtagggcgcccactaacggacaaattgaatccaactttattgcttccgagcacaatgcgccacctcacgacggtatcatgatgggtgaatccccttgctttgcacctggggaagtcccgtacccgcaacaatcttccaaccatgcttatctatcccctggtagtattcccgacatggggggtttccccaacctgacacgaaactacgattacgtagacggcccccaaatattgggaccaggaatggcgcctatcatgaaaccccccaccaacaacgcgataccgaattattacgtcacaagcccacaaatgcatcacaatcaacaccagaaaaacgattgtgacgtcatatccgaaagtagcggtcactcgaacctgagcgatctatcttcaagtccaaggtcatggctgggtgaacttgaccacgtgacacatttccaataactcgttcccctagaaaagttttcactgtgatttctatgccgcccgtacaattattagggttgggttaattattattgaaggtatccgctattgagaaaatatccagagtttttgaccccaacttcccttctccaaccttcaacccccaatttctgacctgttacccgttacctattacccctaactcctattcctaaccccaattgcaaagtttcgcaattccgatgattgagaccaaatattcgcggtaatttgaccccgaaatatctggcaaagtaactttcttctatgcacgcatgactttattgcaaaacggattttcttaaagcgggtgcgttctatgatatatatctcgctaaattctcgtcgtatttgggacttttacacccccattaaagttacaatgggtatttatacagatatctacaaactattacgcatccatccccgctgttaaccaatcaaatcgctctgttatacgcttcggtgacaacgtatttctgtaagacgcattatacgccaatcaagtcgctttaacccctaactaaccgaagcgaaacatagcggtttgattggttgtaaaagggatggttgtgtagtagtctgtatttgaaaaacccttttagtgaaggtatatatgcccactatatatgacgcgtttatcgtcgccacatacccgtataactacagggggttgtgttatagttcccttcattccatctcgcttcaatccgccgctaaaacgcaccctgtgccaaattaatgcggaactgtgtttcaccatgttgcaatattagttgtgtgagtacgcgaataaatattcaaaaaaaaaaaaaaaaaa

>Nkx6 in situ probe template (Stolfi and Levine 2011)

atgtccatggaagaatctggttcttttgcgtttgggaacaacgctgcagttgcgatgagcgcttttcaggcggcggcatctgcaggaatgatggaacataagacaaacaacggaatgcccttctaccactaccctgcatttcaacacaacaaccagaatcaaggtcttcgaaatcaacacaccaacttgcaaaatggaaccccctttggtataaacgacattttaaaccggccaatcgctgcaagcttaactgcgcaagattctactcaccacttgagcagtatcgctggaaatgcccaatcaaacccccggtttacgtcaagtatcggtcacggctctatgatggccgccatgtacttgggaggatctggaattgcaaatgctgtttccacttcgagcggatcaagatatccgaaaccactcgccgaccttcccggtcgcccaccaatatattggcctggtatgatgactgatgattggagagaaaagttagcgatgccaggttcaccttcgaccatcgtgatggaaaagtacggacgaaagaaacatacacgacctactttctcagggcagcagatctttgcgttggaaaaaacctttgaacaaagcaagtacctggctggacctgaacgagctcggttggcatattcacttgcaatgacggagtcgcaagttaaggtttggttccaaaacaggcgaacaaaatggcggaagcgtcacgctgctgagatggcaaccgctaagaaacgacaagatggtgaaatatctctaaaggtaaaggagcgtgaacgcggtgaaagtgaaggacaaagagaatctgtgagtggcgaaaattctacacaacgaccttttaacgatacggatggtatgacggcatcgggtcaaatctttgagaatgcggcagacagcttttccgaatccagcgctgggtcgcgtgaaaattcaagaagatgtagcgagcatgaaggaagtgat

Neurog in situ probe template: N. Satoh gene collection ID R1CiGC29n04

Ebf in situ probe template: N. Satoh gene collection ID R1CiGC02i14

**Dkk orthologs and mutants** NxI/V motifs NxF motifs

>Ciona robusta Dkk1/2/4

MQRRDDPSPPPKKPHRTPEAESTSHGNDLALPSHVSQIKHGQVSGGSISYRCAGNRPGQCGPYNRKQRNHNKIEKRKRRACKKRENKCKRNKIRKCHRSRKCRSHAVKRCRRLKRRCMNNRPLVENPPPKNPNDKPASTPTIIPKREKEKCNNTYECADGFCCAQHGLVKRCKPFLRIGHICPRPRQHHKRSPDSYERCDCEAGLACLASTNRHHKCQRTSLIT

>Cr.Dkk3 [Ciona robusta, tunicate]

MKRLVVFSFILLVNFCSGMYRYRLGGELDRQLSPYTIENDANDREEEEREDMLYKYMMEQLIRDIGKENTFENELQDNDSPPSALEELTRLENTIDDLDKDGRWVDEVETLENEFPPNYHNVTEFRTKIGNQSVHVKESIDKNTLGNEGQSEWLSEEITSKSDVKRDRGPWKCSIAFPCSEDEYCFETQTSSECKECKLEGDQCVDNAECCQEPHANYDHQAMCVFGRCKLSTRPGAIGTICEVPDDCDAGMCCADTPGYARSLCKPNARVGERCSVEPDYNPSEYIVFEDGQYFCPCESDLTCVSRGTNKEDNLQDYNTPRFCEMPRLNLDRASTKDKVQIERDLLAKLFPELVVPHWQQRMETIEDQQFTSKMNPSLHKTTKKAMEENELIM

>Dr.Dkk3a [Danio rerio, zebrafish]

MFLLGFSLCLAVVHGIVPEIPKTDMDIIANMETNAAQEQTMSDVLKEVEELMEDTQHKLEDAVHQMDNETAKSSLHPQNVSSNLQNYSAIETIAGNQTISIGERINKTTDNSTEETNNLSSIQPRDKENIVDHECVIDEDCEKGKYCLYETHSSKCLPCKQLDASCTKDEECCAGQLCVWGQCTINITKGDAGTICQYQTDCKEDFCCAFHKALLFPVCIAKPIERERCIISANHLMELLSWDMDGEGPQEHCPCAGELQCQHRGRGALCLKSQNSSEEELTDTLYSEIDYIV

>Dr.Dkk3b [Danio rerio, zebrafish]

MLKSMILCLCVGLAVGSSVHRGAHLDISDTLEEHVAHGQTTLNEMFREVEKLMEDTQHKLEEAVHQMENETTNSLLNGRDFPDNFHDETTTEIKLGNRTIQLIERINKKTDNKTGKTHFSRTLIQNTERWNEVDHECMIDEDCGDGSFCLYEIVTSKCVPCQTTNMECTKDVECCGDQLCVWGVCAQNKTKGQSGTICQNQNDCSPQHCCAFHKALLFPVCRPKPQEGQGCEREGNQLMEVLLWEDEGPREHCPCAAGLLCQQIQKSSVCVDERHASGEGNED

>Xt.Dkk3 [Xenopus tropicalis, western clawed frog]

MPTFLLLILLLGTGAPTPTRSPSEPPDPSDAPQVDPLFSFREEEASLNDMFREVEELMEDTQSKLQNAVKEMEAEEVLAHRLPGWAQLPNNHNESVTEMGNETIHSQKEMTKNTDNHTGSTLYSETVITSLKNNSKRHQECIVDEDCKSGNYCYFADSEYKCLPCKATEPCTRDGECCEGLCVWGQCAHVTKGEGGTICESQEDCNPGFCCAVHSDLLFPVCTPLPGEGEPCLDPSNKLVDIMNWDVQPAGVLGRCPCSQGLVCQPQSHALVSTCQEPSPDDSKRSDLEVPEVIPPFIGIMPQEGQYYEDGTQLSDGPYASPSEERY

>Ct.Dkk3 [Crotalus tigris, Tiger rattlesnake]

MFLLGSTRLLFLLALLGLVFSAPTLDSKKKEQPQEIVLSFPKDETSLNEMLQEVEELMEDTQYKLTNAVK

EMEAEEDGSKKIMGIEFQKLPANYHNESYTDTKVGNKTIHRHQEIDKVTDNKTGSTSYSETIITSIQEDR

KRNHECIIDEDCEIGKYCAFSDLMYNCHICKSQHTHCSRDVECCGQQLCVWGECMKAVSKGENGTICENQ

HDCNPGLCCAFTKDLLFPICTAFPGEREPCHNPSDGLLNLITWELEPDGALDRCPCAHGLICQKQSHSSE

SLCELSLNKTQSEDKNQLLADELSLLDFVTQDIPGEYEERLIKEVQKELEDEVADSGKLAMASDFFGEEI

>As.Dkk3 [Anolis sagrei, brown anole]

MFQFGSRLLFLPILLGVVNAAPALEGDGEEQPNEEIVVTFPRDEASLNEMFREVEELMEDTQYKLSNAVKEMEAEEEGSKRLAEIDFEKLPANYHNDSHRDTKIGNKTIHTHQKIDKATDDKTGSTSYSETVITSIQEGDNKKNHECIIDEDCVTGKYCEFSGLEYKCRDCKIEHKRCSRDVECCGHQLCVWGECMKTASRGENGTICENQHDCNPGMCCAFSKDLLFPVCTPLAGEREPCYDPSNMLLNLITWELEPDGALDRCPCSHGFICQVQSLSSDSLCERSFNKTQTEDEKPLVEDETSLLDFISEDIPGDYGARLIKEVQEGLEEEAVDSKELDIASDLLFGDEI

>Cm.Dkk3 [Chelonia mydas, green sea turtle]

MLLLLVLPLLLGAACAAPAADGGAELPPGEPEPSFPREEASLNEMFREVEELMEDTQYKLRNAVKEMEAEEEGAKNPSEVDFENLPPSYHNESNTDTKIGNKTIHTHQEIDKVTDNKTGSAIYSETVITSIKDGESKRNHECIIDEDCETGKYCQFSSFEYLCQLCKTQHAPCSRDVECCGDQLCVWGECVEAASKGKTGTICENQHDCNPGTCCAFQKDLLFPVCTPLPAEGKPCHDPLNKLLNLITWELEPDGVLERCPCASGLICQTHSHSTVPVCEVSFNDTGSNEKEDALLMDEISFLGLIPRDVLGDYEDSSIIQAVRKELESLEENTSEQTDLKEPDLAHDLLFGDEI

>Am.Dkk3 [Alligator mississippiensis, American alligator]

MPGSRRRRLLLLLSLLLLLGAAWAAPAGAEEPSFPREEASLSEMFREVEELMEDTQHKLRHAVQEMEAEEEGSKKVSEDDFENLPPSYHNESNTETRIGNKTIQIHQEIDKVIDNKTGSAVYSETVITSIKDGEIKRNHECIIDEDCETGKYCQFSSFEYKCQPCKAQHTHCSRDVECCGEQLCVWGECTQAVSKGENGTICENQQDCNPGMCCAFQKDLLFPVCTPLPAAGEPCYDPSNRLLNLITWELEPDGVLERCPCASGLICQPQSQNNMPTCELSFNETKSNDKDDPLLMEEIPFLGFMPRDVLGDYDDSSIIQEVRKELESLEENISEQTDIKEPDLAHGLLLGDEI

>Tg.Dkk3 [Taeniopygia guttata, zebrafinch]

MLREVEALMEDTQHKLRNAVQEMEAEEEGAKKLLEVSFEDLPSNYHNESNTETRLGNRTVETHQEIDKVTDNKTGSTVFSETVVTSIRDGENKRNHECIIDEDCEPGKYCQFSTFEYKCQLCKPQHTPCSRDVECCGEQLCVWGQCRRSISRGENGTICENQHDCNPGTCCAFHKELLFPVCTPLPEEGEPCHDPSNRLLNLITWDMEPDGVLEQCPCAGGLSCQPQSHRTTPVCQLSANETQHTEKEDPLIMDEMPFLSLLPQDLLSDYEESSVIQEVRRELESLEDQAGLKPEPEAVQELVLGDEI

>Gc.Dkk3 [Gymnogyps californianus, California condor]

MRRGAGPGPRRRWLLLAALLGSLCCAAAGGGGRRRAASLGEMLREVEALMEDTQHKLRNAVQEMEAEEEGAKKLSEVNFENLPPNYHNESNTETRIGNKTVQTHQEIDKVTDNKTGSTVFSETVITSIKDGENKRNHECIIDEDCETGKYCQFSTFEYKCQLCKTQHTHCSRDVECCGDQLCVWGECRKSTSKGENGTICENQHDCNPGMCCAFQKELLFPVCTPLPEEGEPCHDPSNRLLNLITWELEPDGVLERCPCASGLICQPQSHSTTSVCELSANETRNNEKEDPLIMDEMPFLSLLPRDILSDYEESSVIQEVRKELESLEDQAGLKPEPDSAHNLFLGDEI

>Mm.Dkk3 [Mus musculus, mouse]

MAKSGFSWLGGGQRGTNMQRLGGILLCTLLAAAVPTAPAPSPTVTWTPAEPGPALNYPQEEATLNEMFREVEELMEDTQHKLRSAVEEMEAEEAAAKTSSEVNLASLPPNYHNETSTETRVGNNTVHVHQEVHKITNNQSGQVVFSETVITSVGDEEGKRSHECIIDEDCGPTRYCQFSSFKYTCQPCRDQQMLCTRDSECCGDQLCAWGHCTQKATKGGNGTICDNQRDCQPGLCCAFQRGLLFPVCTPLPVEGELCHDPTSQLLDLITWELEPEGALDRCPCASGLLCQPHSHSLVYMCKPAFVGSHDHSEESQLPREAPDEYEDVGFIGEVRQELEDLERSLAQEMAFEGPAPVESLGGEEEI

>Hs.Dkk3 [Homo sapiens, human]

MQRLGATLLCLLLAAAVPTAPAPAPTATSAPVKPGPALSYPQEEATLNEMFREVEELMEDTQHKLRSAVEEMEAEEAAAKASSEVNLANLPPSYHNETNTDTKVGNNTIHVHREIHKITNNQTGQMVFSETVITSVGDEEGRRSHECIIDEDCGPSMYCQFASFQYTCQPCRGQRMLCTRDSECCGDQLCVWGHCTKMATRGSNGTICDNQRDCQPGLCCAFQRGLLFPVCTPLPVEGELCHDPASRLLDLITWELEPDGALDRCPCASGLLCQPHSHSLVYVCKPTFVGSRDQDGEILLPREVPDEYEVGSFMEEVRQELEDLERSLTEEMALREPAAAAAALLGGEEI

>Cl.Dkk3 [Clavelina lepadiformis, tunicate]

MMKSIILNRYIHVYCCVKDITKMVGVLLNLAFCGLMIETALCSPYSRLAGSSSSSNGLYNNNDNGFYDIPMEEDVQNQLINEETKLLQDLLVSDLLEDLKEVGAADGLDNNLVGSPVDPLIELNDLEDAFIEEEDDDGKWLDDVQEVINSLPANYHNSSEYQTQIGNENVDVQQTINKETDDIGNEMSLSDQMQVQTKDGNGAFACSLANPCNEDEYCYKTDMYSECLVCLEVGEQCIDDAQCCSDNDDDLAHCVAGACAAGVKSGTVDTICEVAEDCDEGLCCATVQGYSRKMCKRFARLNERCSVETSVKTDVYIASINYYCPCVEGLRCQGRKPPVYNFLPVRYRDGQFCVLDPGSIIDKFRPSSRQNNQDMLDSQPIDQDLLDLEARLENENDQRNLNQDSSQQHHNAKKAMEMEPVEFMM

>Ciona robusta Dkk3 NxI mutant

MKRLVVFSFILLVNFCSGMYRYRLGGELDRQLSPYTIENDANDREEEEREDMLYKYMMEQLIRDIGKENTFENELQDNDSPPSALEELTRLEATADDLDKDGRWVDEVETLENEFPPNYHNVTEFRTKIGNQSVHVKESIDKNTLGNEGQSEWLSEEITSKSDVKRDRGPWKCSIAFPCSEDEYCFETQTSSECKECKLEGDQCVDNAECCQEPHANYDHQAMCVFGRCKLSTRPGAIGTICEVPDDCDAGMCCADTPGYARSLCKPNARVGERCSVEPDYNPSEYIVFEDGQYFCPCESDLTCVSRGTNKEDNLQDYNTPRFCEMPRLNLDRASTKDKVQIERDLLAKLFPELVVPHWQQRMETIEDQQFTSKMNPSLHKTTKKAMEENELIM

>Ciona robusta Dkk3 NxI/F mutant

MKRLVVFSFILLVNFCSGMYRYRLGGELDRQLSPYTIENDANDREEEEREDMLYKYMMEQLIRDIGKEATAENELQDNDSPPSALEELTRLEATADDLDKDGRWVDEVETLEAEAPPNYHNVTEFRTKIGNQSVHVKESIDKNTLGNEGQSEWLSEEITSKSDVKRDRGPWKCSIAFPCSEDEYCFETQTSSECKECKLEGDQCVDNAECCQEPHANYDHQAMCVFGRCKLSTRPGAIGTICEVPDDCDAGMCCADTPGYARSLCKPNARVGERCSVEPDYNPSEYIVFEDGQYFCPCESDLTCVSRGTNKEDNLQDYNTPRFCEMPRLNLDRASTKDKVQIERDLLAKLFPELVVPHWQQRMETIEDQQFTSKMNPSLHKTTKKAMEENELIM

**VAChT promoter (-2083/+3) alignment and TF binding sites**

CLUSTAL format alignment by MAFFT (v7.511)

(See Supplemental Table for binding site scores by JASPAR)

**Potential Neurog sites**

**Potential Onecut sites**

**Unknown repeated sites: splicing-related stem loop? See Mathews et al. 2015**

(exon 1, 5’ UTR in robusta, unknown if conserved in savignyi)

(exon 2 in robusta, unknown if conserved in savignyi)

VAChT start codon

Savignyi	------------------------------------------------------------

Robusta	attacgtcgtaaacctttggctaccat**catctg**cctcaaaacaaaattaattaaagaaat

Savignyi	--------------------ttaaaagcaaccaa--------------------------

Robusta	gcgttagtgtatcctttgactcggaatcaatcaaatcaag**caaaaatcaatatgtg**aaat

*...** ***.***

Savignyi	----------------------------------------------------catttcgt

Robusta	taaccatttagaccttgtgtcattcccattgcggtgacttgtcctagttgtgcgttttta

*.***.

Savignyi	tcgtaaaactcgtctttctttcattaaaatcaacttgga**catttg**aattttgaattaaaa

Robusta	tcagcagattgatttaacagtcaccggaacaggcaagacaatatttcaaaccagccaatg

**. *.*.* .*.* * ***....**. ..* *. ** *	. *...** .

Savignyi	ttccgatcgtttttttaatgcaaaaaagcaactaaatctattttaaaatcgtatttttca

Robusta	ttaatttcaga**caaatg**aagcaatctgaaaatcagaacaaatcaaaaaacataagttttg

**	**. . *.* ****	.. **..*.* * * *. **** *.** ***..

Savignyi	attccaaaattatctaattatagcgcaagttaa**catatg**gctttaaagagcaatagggtt

Robusta	gtttttaaagcatagaa---------aacgtaccgtattttttaatgtaattgttaaatt

.**.. *** .** **	** ** *.*** .** * . *.. .* ...**

Savignyi	acgtactctatcgtctgaggtttttggcgtcttataaataaaaaaatacaatttttatta

Robusta	ttgttatttaatatagtagggt---------------agggggagatgggacactttttc

.** *.** ..*	*** *	* ....*.**. .*. .** **

Savignyi	actgcattcatttaaatttccctgcattattattatgttaaaaatggatataaaaagatc

Robusta	attctgttttctcgtcttggtagcaaacaaaaacattaaaagaat---tataaaaccgta

*.* ..**. .*.. ** . * .* * .**	**.***	******* .*

Savignyi	ttagttcggtatttgtaaattctgcagaacaattattactggtgacgtattgtaaacgta

Robusta	tcctcgcgactcctacagaccgtttaaaacaggatatttggatattctgttctaaaggtg

*. . **.. ..*..*.*.. * .*.****. * . *.*. . *.** **** **.

Savignyi	aggcttcctttttcgccgctatctcccct**aaaattgatttttc**tcaaaatttctggacca

Robusta	tcccatcttaccccacagtactatacacta--------------tacacattctgtaccg

* **.* ...*.* *. * * * ***	.* * ***** ***.

Savignyi	atgtactttagcctttcgctaattagacattcgggtacagggccgaac----actataac

Robusta	atatatttta--------ttgattaaatttgaagttgttaaacttaattacgattaaatt

**.**.****	.*.****.*. * .* *.. ...*. **. *.** * .

Savignyi	tcgacgcacgaaaaaggcaactcgacgttagctccattgtctatcgcttcacgtcgacgg

Robusta	tcggcaaattgaaaatgagccatgaattaatcaaaaattattcttgatcgttgttttgta

***.*. *. .**** * . * .** * * *	* * .* *.* *. .**. .

Savignyi	ggatggtagtcc**aacattgattttat**cgcttgagctctatatcgggctcgggttgttaaa

Robusta	aaacataact**tttttttgattttttt**gggaggcgccctgtgttccactt--attattatt

..*.. * *.. . * . **** * *	* **.**.*.*. .**. .**.***

Savignyi	gatttgatcaatcgcgctaagcaataccggcaacgccaaagtcgttttttttttctaaag

Robusta	tgtttgttttgctatgtcaa-----------------------tactttaatataaaaga

.**** *. .....*..**	.*** * * **..

Savignyi	cattttgaccgaactgtgttgcattacagcaccagactcagatttggttgtgaaagtgac

Robusta	aattaatattggctgacatttcaatttaacgctaggcttatgttttgtttcgtgaataat

***	*..*. . ...** ** * .*.*.*.**.**.* .*** *** .* .*.*.*.

Savignyi	atatat------gatcggttatttgcaggtttttatattttaatagttttatataaagcg

Robusta	ctgcataaagaaaaaaagcagtatcgactctcctattgttgaatcactct-tgcttccct

*..**	.* .*. .* * * .*..*** ** *** ..*.* *.. *

Savignyi	gtcatagtgttaggcaactttgagtcgttaaagtcgtttttttcacgttttgtgcatcct

Robusta	tccattgtgcaggagatagtgacgcaatg**aacattgatttt**aacgcttctagt-------

.*** ***. .*. *	* . *. .* ** .*.* **** *.* *.* **

Savignyi	tactagacctgtagtgcattagaccatatcttaatgattaaaataaacacagttatttag

Robusta	--------**cagttg**ggcccttgctca**ca**----**attg**cctaaaatttgat**caattg**tggat

* ** * ** .* * .**.*	* ** .****** . **.**.* *

Savignyi	tgtaacattgatacgaact-----attgagatacttataagttcataaaaattaagacgc

Robusta	tgaaaagttaatattctttcctgaattaacatgtctataggt---aaggtattgagacgg

** ** .**.***. .*	***.* **...****.**	*.. ***.*****

Savignyi	tggttatgtattttttt-----ctgttttaccacattttcttatcttaggttggtagaat

Robusta	ctcccatttaatttcttgtccgatctttataaaaaaatatttgttaaaggtacgtttaat

. ..** ** ***.**	* ***	* * * .**.*. **** ** ***

Savignyi	ttagttatttcaatcataacgcctgattttgattgtgaagaagtaaaatccggacaagca

Robusta	ttgaacatatagtgcagaagtcttcggaattttcatatagagttattttttagata--ct

**.. .** * . ** ** *.* . * *..*. ***. **	*...**.* *

Savignyi	atttgcgcgttaaatcggtaagtaagcgttgcgtattttcacggggtggatatattcgtg

Robusta	atatgcgacatattttcatagcccaaattaaattgttttataatttgtttggcttttaca

** ****	** *. .**. . *. * . *.****	. .. **....

Savignyi	tattttagcatatttaaccaatatggtt--atcgtaccccctccctgtaaaaaaacaaag

Robusta	gattttaatttaacttaggtgtacgattcaatagaaagattttaacctgtaaaaatacgt

******.. ** .* *	.**.*.** ** * *	..*. . *. *****.* .

Savignyi	ca**aaattcattttt**gtgtccatttatat**catttg**attaccacatatagggtgga**cagat**-

Robusta	gcaagt---tgtttggaaataccttttttgttggacaaagttgtatcaggtaagttgata

**.*	* **** . .*..* * *..** **. *	..*** .***.... ***

Savignyi	--**g**tatgttatacaacaacttttgtatttgaatttcgtttccgtatta----attatgca

Robusta	gcatgtatcttaaatctggcgtggtgtttatgtattggcttcatcatgaaatagtttgtt

.*.*.*. ** * * . . * **.***. .* *.* .*.*.* *. * * **.

Savignyi	tagaaggttttgtgtgcaacgctatttctgatgaattgggtgccacactgtgacccgaca

Robusta	ttgtccttttacttgtcaattttatttaaattacatgagggtacaattcattataacgta

* *	*** *	***. .***** . *. ** .**	** ...* *. ..*

Savignyi	cgc-----aagtaccaccgacgacttaccttgtccaggttatagcttagagggaagtcga

Robusta	cttcggtgaaaggttaatgttaa**caaatg**ccgtccggtcttctgcgtatctttcgatttc

* . **. ...* .* ..** *. ..****.* .* . ** **	..*.

Savignyi	gatgcactttggaagcggattttttatag**ttttaatcaataaa**gcgcgacgtgcatgaaa

Robusta	gtattaattatgcatagaaagggtttaagtgcaatgctatttctgatgatgtgtgtgtaa

*	.* ** * * *.*	** *** . * * **	..**.***..** **

Savignyi	cca**cacgtg**ggtcgggtgtctgaatacgctttgcgtggtcttgtggattcggtaactttg

Robusta	cacca--acgacctgatgacgaaaaac----ttgactgtttattaaataaaatagcgcaa

* **	*..* *.** * .** **	* .. **.* *..** ..**.* . .

Savignyi	aatggaaaaggcagtcaattttcatacctttctagtgatgagcccatcatg--ggttcga

Robusta	caagacggcggaatttataaatggcatgttgctcgtaaacagcagcgcgtgcctgtttta

* *. .. ** * *.*	* ..*. ** ** **.* ***	*.**	***. *

Savignyi	aacttgggccggttaaaaaaagaaatttgttttcagaattataaacggttgtacggccga

Robusta	actctggagaaacaagttatattcaactatttgttatctcacaaagcacggtattgcact

* ..***. ... *. * *	* .*.*** . . *.*.*** .. ***. **

Savignyi	atggttaacgcgaacgatatagttaccgatatgtacacaaaaatatcatagtaacaggcg

Robusta	cttatctaatgtactaatacaaatatatatatatatatatatatatatatctagctgttg

* .*. *	* ..***.*. **. ****.**.*.* * ****	**.* * .*

Savignyi	tataggttcgaagcttgggctggtaaaaacgcattgtaatatccaaaacgcataccgaat

Robusta	tcgaggtaagatgaactgcagttaacaataaaaaccaacttcgttatacacaaattcaat

* **** ** * . *	* ** . * . * * . . * **.** *.. ***

Savignyi	agatactgtgtgtgaaatagcttagaaaaccggtatttcgttttatttcta**caaaaggca**

Robusta	ataaagcgacagctcagtgttgtagagtaggtacccaactttcttaatctg**caaaaggca**

* * * .*	*. *.*. . ****. *	.. . * **.*	***.*********

Savignyi	**aacacatg**gcttagtaactacgttaactaaaataataatgttactaaactgtc**tttgcct**

Robusta	**aa**tacatgattta--taatatgtaactcagcataacaac---------ctgtt**tttgcct**

**.*****..***	* **.** * ..*. ****.**. ****.*******

Savignyi	**tttg**cagat--cgacgaaagaccggtggtctatactcga--------aagggaagatttt

Robusta	**tttg**cagattacatctgagggcgggttgtgtgggtcagaaatttatcaaaggaaaataaa

********* *. * .*.*.* *** ** *. ... **	**.****.**

Savignyi	aaagattaaa---aaagttgcttctactgagaataccaatatggaaatgttgaaattatt

Robusta	gttgattgaaagcaaatttgtttctact------------------------ttattgtt

. ****.**	*** ***.*******	***.**

Savignyi	cggcaatatgacaatgacatttctgcatcgctgcgttgacggaatg

Robusta	catc---------------------------------------atg

*. *	***
